# Genetic ancestry influences the relative expression of ancestral haplotypes within admixed individuals

**DOI:** 10.64898/2026.11.11.765740

**Authors:** Esteban Vazquez-Hidalgo, Makaela Mews, Lissette Gomez, Tianjie Gu, Anthony Griswold, William S. Bush

## Abstract

Allele-specific expression (ASE) reflects differences between the two haplotypes a person carries, but most ASE studies have been done in European-descent samples. Admixed individuals carry haplotypes from different ancestral backgrounds within the same cell, which allows ancestry effects on regulation to be measured directly. We integrated phased whole-blood RNA-seq with local ancestry inference in 779 participants from the MAGENTA cohort, including non-Hispanic White, African American, Caribbean Hispanic, and Peruvian individuals. Higher West African (YRI) ancestry proportion was associated with more ASE-positive genes per person, and the association remained after adjusting for age, sex, Alzheimer’s disease status, APOE genotype, and the number of heterozygous genes. Peruvian ancestry showed no significant association. A gene-level test of imbalance across population groups found 2,053 of 15,008 genes with significant differences, including the major histocompatibility complex in all four groups. A genome-wide scan of 2,213 1 Mb windows compared people carrying two haplotypes of different ancestries with those carrying two of the same ancestry. It identified 52 non-redundant loci that replicated across all four population groups, containing 201, 60, and 110 ancestry-associated genes under the CEU, YRI, and PEL reference baselines. A haplotype-resolved directional analysis showed no genome-wide advantage for either ancestry. It found locus-specific dominance at 11 loci, and only chr8:2-3 Mb (CSMD1) showed the same direction in more than one population group. Genetic ancestry therefore shapes allelic regulation at global, gene, and locus scales.

## Introduction

Allele-specific expression (ASE) analysis provides an assessment of the relative expression of the two chromosomal copies of each gene a person carries (Dimas et al., 2009; Castel et al., 2015; Aguet et al., 2017) and can be driven by multiple potential mechanisms. Genetics of the two haplotypes may differ, with collections of cis-regulatory genetic variants driving differences in gene expression, or epigenetics may differ, with the two haplotypes having different chromatin accessibility, histone modifications, or DNA methylation profiles (Kasowski et al., 2013; Aguet et al., 2020; Dimas et al., 2009). While the bulk of disease associated variants are regulatory, prominent examples of common disease-associated coding variants (such as *APOE* or *APOL1*) account for a substantial proportion of disease risk. Understanding ASE will advance our ability to model and interpret genetic risk.

Much like studies of expression QTLs (eQTLs), examinations of ASE have mostly been conducted in European-descent populations (Aguet et al., 2020; Stranger et al., 2007; Lappalainen et al., 2013; Bhattacharya et al., 2022). Widely used resources such as GTEx provide deep tissue coverage but are predominantly of European ancestry, reducing power to model ancestry-diverse regulatory mechanisms. Prior studies have shown that expression differs by genetic ancestry across individuals (Mogil et al., 2018; Gay et al., 2020; Kachuri et al., 2023), and that both global and local ancestry contribute to expression variation in admixed populations (Price et al., 2008). However, large-scale expression studies commonly adjust for global ancestry alone (Mogil et al., 2018), which could mask local ancestry effects that occur at the locus level (Zhong et al., 2019; Gay et al., 2020). Whether the two haplotypes of an admixed individual are expressed at different levels depending on their ancestry has not been tested genome-wide across multiple populations. Whereas global ancestry describes the genome-wide proportion of alleles derived from each reference population, local ancestry assigns ancestral origin of chromosomal segments, enabling detection of regulatory effects that are haplotype-specific rather than proportional across the genome. This distinction is critical for ASE, which is measured within individuals at specific heterozygous sites where the two haplotypes may derive from different ancestral backgrounds. Studies of admixed populations are particularly well positioned to examine differences in expression by ancestry, as they naturally carry haplotypes from multiple genetic backgrounds within the same individual (Bhattacharya et al., 2022).

Genetically admixed individuals provide a distinct opportunity to explore ASE as the two copies of a gene may have been inherited from different ancestral backgrounds. For example, Hispanic/Latino groups exhibit complex three-way admixture involving European, African, and Amerindian components (Bryc et al., 2010; Granot-Hershkovitz et al., 2022), allowing comparisons of multiple different ancestral versions of a gene, each shaped by demographic history and local adaptation leading to distinct regulatory architectures. A growing number of admixture mapping studies have demonstrated that local ancestry is a potential modifier of genetic effects, including eQTL signals and disease-associated loci (Zhong et al., 2019; Gay et al., 2020; Natri et al., 2024). Despite these examples, it remains unclear if the ancestral origin of a region consistently modifies regulatory dosage of genes, and how such effects vary across populations, a gap that limits how well admixture mapping signals and risk prediction models can be interpreted across ancestry groups. Any consistent bias in expression of one ancestral copy over another may be useful for constructing ancestry-aware risk prediction models and may point to specific regulatory effects within a particular ancestry group.

To address these gaps, our study integrates allele-specific expression with local ancestry information to characterize ancestry-dependent regulatory variation in an admixed cohort. The key methodological opportunity in admixed cohorts is that admixture creates within-individual ancestral contrasts at heterozygous loci: an individual with European and West-African ancestry carries, at many loci, one haplotype shaped by European regulatory history and one shaped by African regulatory history.

Integrating local ancestry and ASE at the gene level allows us to examine both common and ancestry-enriched allelic effects, revealing regulatory variation that may be overlooked by population-level eQTL studies (Heap et al., 2009; Moyerbrailean et al., 2016; Knowles et al., 2017; Nuytemans et al., 2020; Griswold et al., 2020). This within-individual design allows us to attribute allele-specific expression to a specific ancestral background, a natural experiment that is unavailable in genetically homogeneous cohorts. By systematically mapping ancestry-ASE relationships across thousands of genes, we will establish the relative expression of genes based on their ancestral background which can inform risk prediction models in admixed populations and provide biological interpretations of local ancestry association signals.

## Methods

### Study Cohort

This study analyzed data from participants in the Multi-Ancestry Genomics, Epigenomics, and Transcriptomics of Alzheimer’s Disease (MAGENTA) project, which leverages samples collected across multiple ongoing studies. Participants were recruited at sites in the United States, Puerto Rico, and Peru and included non-Hispanic White (NHW, n = 210), African American (AA, n = 193), Caribbean Hispanic (CHISP, n = 293), and Peruvian (PER, n = 83) individuals, for a total of 779 participants. All participants or their legally authorized representatives provided written informed consent under protocols approved by site-specific Institutional Review Boards. Although the overall sample size is modest relative to large biobank studies, MAGENTA was specifically designed to study ancestrally diverse admixed populations, the very individuals required to understand local ancestry effects on gene regulation, making it uniquely suited to the questions addressed here despite the smaller cohort size.

### RNA Sequencing Data Processing

Whole-blood samples were collected at the time of clinical evaluation for Alzheimer’s Disease. Total RNA was extracted and subjected to high-throughput paired-end RNA sequencing following standardized protocols (Griswold et al., 2020). Briefly, RNA was extracted from frozen whole-blood samples, quality controlled with RNA integrity (RIN) scores ≥5, prepared using the NuGEN Universal Plus mRNA-seq with globin and ribosomal depletion, and sequenced using 125-bp paired-end Illumina HiSeq3000. RNA-seq FASTQ files were processed using the STAR aligner (v.2.7.10a) (Dobin et al., 2013) and aligned to the human reference genome (GRCh38) in a splice-aware manner. Reads were quality controlled using Picard, with samples failing RNA-seq quality metrics (i.e. low sequencing depth, evidence of mapping bias, etc.) were excluded from downstream analyses. Gene annotations were derived from GENCODE v43 (Harrow et al., 2012).

### Genotype Data Processing

Genome-wide array-based genotype data were obtained for all individuals. Standard quality control procedures were applied, including filtering of variants based on call rate, Hardy–Weinberg equilibrium, and minor allele frequency, as well as exclusion of individuals with excessive missingness or discordant sex information. Genotypes were aligned to the GRCh38 reference and harmonized across cohorts. Genotype phasing and imputation was performed using a reference-based approach (Browning and Browning, 2016) via submission to the NHLBI TOPMed (v2) imputation server, yielding phased haplotypes across autosomal chromosomes. Genotypes were filtered using an R^2^>0.3, which retains some lower quality genotype calls that are ultimately resolved via comparison to variants within the RNA sequencing reads. Phased genotypes were required for downstream allele-specific expression and local ancestry analyses (Taliun et al., 2021).

### Allele-Specific Expression Quantification

Allele-specific expression was quantified by integrating phased genotype data with RNA sequencing reads overlapping heterozygous variants. Heterozygous single-nucleotide variants were identified from high-quality, phased genotype data and intersected with aligned RNA-seq reads to obtain allelic read counts. Analyses were restricted to autosomal loci, and only uniquely mapping reads with sufficient base and mapping quality were retained. To minimize technical artifacts, allelic read counts were generated following established best practices for ASE analysis, including filtering of sites prone to allelic mapping bias and exclusion of reads that could introduce double counting due to overlapping mate pairs or PCR duplicates. These procedures reduce false signals of allelic imbalance arising from alignment artifacts, genotyping error, or technical variability.

Haplotype-aware ASE quantification was performed using a read-backed phasing approach implemented in phASER, which assigns RNA-seq reads to phased haplotypes based on overlapping heterozygous variants (Castel et al, 2016). This strategy enables aggregation of allelic expression across multiple sites within a gene while preserving haplotype consistency, thereby improving robustness and interpretability of ASE estimates. Allelic read counts were summarized per haplotype for each gene and individual, yielding a quantitative measure of allelic imbalance that reflects cis-regulatory dosage effects (Castel et al., 2016).

Sites and genes with insufficient read coverage or unreliable haplotype assignment were excluded from downstream analyses. Allelic imbalance was analyzed as a continuous quantitative measure in the ASED analyses, enabling comparison of regulatory dosage across genes and individuals, and as a binary ASE-positive classification when counting ASE-positive genes per individual.

To summarize regulatory imbalance at the gene level, phASER aggregated allelic read counts across all covered heterozygous sites within each gene for each individual, yielding a read count for each of the two haplotypes (that is, the reads mapping to each haplotype across the gene body). Following phASER recommendations (Castel et al., 2015), genes with at least 8 total reads in an individual were tested for ASE using an exact two-sided binomial test of the two haplotype-specific read counts against the null hypothesis of equal allelic expression, and observations below this read threshold were not tested. P-values were adjusted by the Benjamini-Hochberg procedure, and gene-individual observations with a false discovery rate (FDR) q-value ≤ 0.05 were classified as ASE-positive. No minimum fold-change threshold was applied. Genes with all reads from a single haplotype (monoallelic) were classified as ASE-positive. Genes without sufficient coverage or reliable haplotype assignment were excluded from ASE calling.

This framework produces a continuous gene-level measure of regulatory dosage that reflects cis-acting allelic imbalance within individuals, enabling comparison of regulatory effects across genes and samples.

### Local Ancestry Inference

Local ancestry was inferred along phased haplotypes using a reference-based ancestry deconvolution approach in FLARE (Browning et al., 2023). Reference panels from the 1000 genomes high coverage release representing major continental ancestries were used to assign ancestry states along each chromosome, and local ancestry calls were harmonized with phased genotype data (Byrska-Bishop et al., 2022). For each variant site and individual, the ancestral origin of each haplotype was recorded and integrated with allele-specific read count data to enable ancestry-stratified analyses of cis-regulatory imbalance.

For each gene and individual, the ancestry of each expressed haplotype was determined by the majority local ancestry assignment across heterozygous sites within the gene on that haplotype, as inferred from phased RNA-seq reads assigned by phASER. For each individual at each locus, we define local ancestry zygosity (LAZ) as the combination of local ancestries on the two haplotypes, whether both derive from the same continental ancestry (LAZ-homozygous, e.g., CEU/CEU) or from different ancestries (LAZ-heterozygous, e.g., CEU/YRI).

### Statistical Analysis

Distributions of gene-level allelic imbalance were characterized across individuals and genes. Robustness and reproducibility of gene-level ASE measures were assessed across technical conditions and subsets of samples. Downstream analyses evaluated relationships between gene-level regulatory dosage, genetic context, and ancestry background using regression-based models adjusted for relevant covariates, including age and sex where appropriate. Multiple testing correction was applied where applicable. All statistical analyses were performed using R version 4.3.1 (R Core Team, 2023).

### Global Ancestry and ASE Burden

To characterize inter-individual variation in ASE burden, quasi-Poisson generalized linear models (GLMs) were fit using per-individual gene counts as the dependent variable evaluated across three ASE-positive partitions: all ASE-positive genes, ASE-positive protein-coding genes, and ASE-positive non-protein-coding genes, with non-protein-coding counts calculated as the all-gene count minus the protein-coding count. The same models were also fit to heterozygous gene counts in the three matching partitions, and these results are reported as a supplementary analysis. Models were fit as:

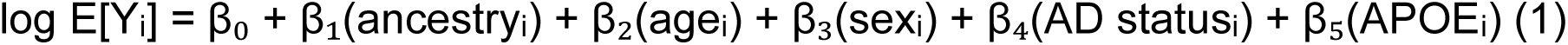

where Yᵢ is the per-individual gene count outcome, ancestryᵢ is the continuous global ancestry proportion (YRI or PEL, each tested in a separate model), and the variance of Yᵢ is proportional to its mean, Var(Yᵢ) = φE[Yᵢ], with the dispersion φ estimated from the data.

A sixth covariate model additionally included the heterozygous gene count of the matching partition as a predictor (ASE-positive outcomes only) to assess whether the ancestry effect on ASE burden was attributable to admixed individuals carrying more testable heterozygous sites. The model was specified as:

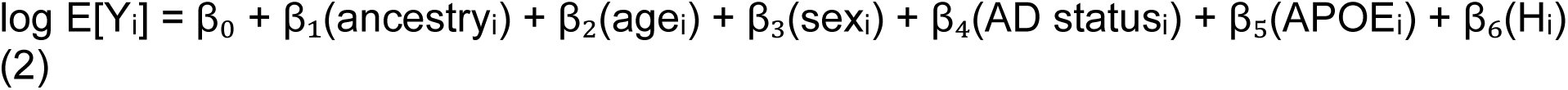

Here, β₆ is the coefficient for Hᵢ, the number of heterozygous genes in the matching partition carried by individual i (all genes for all-gene ASE counts, protein-coding genes for protein-coding ASE counts), and all other terms are as defined above. Heterozygous partitions capture genes with *potential* to exhibit allelic imbalance while the ASE partitions capture genes that do. Continuous global ancestry proportions (YRI and PEL components), age, sex, Alzheimer’s disease status, and APOE genotype were included as predictors, where AD status was coded as AD versus cognitively unimpaired (which includes MCI and other diagnoses) and APOE genotype was coded as a categorical variable with 3/3 as the reference. Linear and quadratic ancestry models were compared by likelihood ratio test to assess evidence of non-linearity. To quantify the variance in ASE-positive protein-coding gene counts attributable to ancestry, we compared the pseudo-R² of three models: ancestry alone, age, sex, AD status, and APOE genotype without ancestry, and the fully adjusted model. Pseudo-R² was calculated as 1 − (residual deviance / null deviance), and for the heterozygous-count-adjusted model we also compared the pseudo-R² with and without YRI ancestry (Supplementary Table S2). To assess confounding, we compared the YRI coefficient across the sequential covariate models.

### Gene-Level Population Differences in Allele-Specific Expression

To identify genes exhibiting heterogeneity in allelic imbalance across population groups, a gene-level adaptation of the ASED framework (Tian et al., 2018) was applied. For each gene, per-sample allelic imbalance (AI) was computed as the absolute difference between allele-specific counts divided by total read coverage:

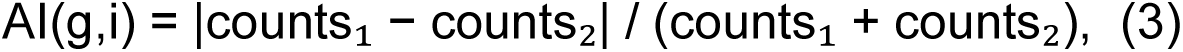

where counts₁ and counts₂ are the haplotype-specific read counts summed across heterozygous sites within the gene body for that individual. A Kruskal-Wallis rank statistic with permutation-based p-values was used in a two-stage testing procedure (a 500-permutation screening pass and a 2,000-permutation refinement pass) to test for distributional differences in imbalance across self-identified cohorts (NHW, AA, CHISP, PER) at a 5% FDR threshold, with Benjamini-Hochberg correction across all 15,008 tested genes, and genes were reported as significant only if at least 20 individuals were informative in each of the four groups.

### Local Ancestry-Stratified ASE

To isolate the contribution of local ancestry to allelic imbalance, local ancestry calls were used to assign the ancestral origin of ASE-informative variants at each locus, enabling stratification of ASE by local ancestry state. Genes with sufficient within-individual variation in local ancestry states were tested for association between ancestry state and allelic imbalance at a 5% FDR threshold.

### Ancestry-ASE Scan Overview

To identify genomic regions in which the local ancestry configuration of an individual’s haplotypes is systematically associated with the probability of allelic imbalance, we developed an ancestry-ASE scan method. At each locus, an individual’s diploid local ancestry zygosity (LAZ) state describes the ancestral origin of the two haplotypes carried at that position. Possible LAZ states include homozygous configurations (CEU/CEU, YRI/YRI, PEL/PEL), in which both haplotypes derive from the same reference ancestry, and heterozygous configurations (CEU/YRI, CEU/PEL, YRI/PEL), in which the two haplotypes derive from different ancestral backgrounds. The scan tests whether ASE at heterozygous variants within a genomic window is associated with the local ancestry pair carried at that locus, using a non-overlapping window logistic regression framework with cross-population replication as the primary significance criterion. Within each 1 Mb window, a binomial logistic regression was fit:

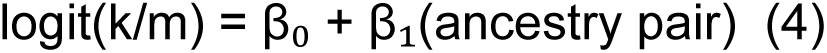

where *k* is the number of ASE-positive variants and *m* is the total number of LAZ-heterozygous variants carrying a given ancestry pair configuration within the window. The ancestry pair term contrasts each heterozygous configuration (e.g. CEU/YRI) against a homozygous reference baseline (CEU/CEU, YRI/YRI, or PEL/PEL). The log odds ratio (β₁) and its 95% confidence interval were extracted for each window and ancestry pair contrast.

The genome was partitioned into non-overlapping 1 Mb windows. Within each window, we fit a logistic regression model separately within each admixed population (AA, CHISP, NHW, PER) with ASE status as the binary outcome:

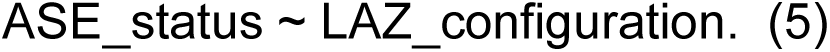

LAZ_configuration is a categorical variable encoding the diploid local ancestry state at that window, with a LAZ-homozygous configuration specified as the reference level (CEU/CEU, YRI/YRI, or PEL/PEL depending on the baseline scan). This design tests whether carrying a mixed-ancestry haplotype pair (LAZ-heterozygous) increases the probability of allelic imbalance relative to carrying two haplotypes from the same reference ancestry. The log-odds coefficient on the LAZ-heterozygous term is the primary test statistic. Windows with fewer than 200 total variant-level observations (heterozygous variant–individual pairs) were excluded to ensure stable model estimates. P-values were adjusted genome-wide within each population using the Benjamini-Hochberg procedure. ASE status was defined at the variant level: a heterozygous variant in a given individual was considered ASE-positive if it was covered by at least 20 reads, passed the binomial test for allelic imbalance at q ≤ 0.05, and showed at least a two-fold difference between haplotypes (|log₂ allelic fold change| ≥ 1.0). This variant-level definition applies only to the ancestry-ASE scan and is separate from the gene-level ASE call described in the Methods. Variants were assigned to windows by genomic position. A schematic overview of the full scan pipeline is provided in Figure 6.

Three independent scans were conducted using different LAZ-homozygous reference baselines: CEU/CEU, YRI/YRI, and PEL/PEL. Running three separate scans, each with a different homozygous reference group, tests whether LAZ-heterozygous configurations show elevated ASE relative to each ancestral background independently. A window was retained as a candidate locus only if it reached q < 0.05 in all four population groups within the same baseline (cross-population replication criterion). This replication requirement ensures that identified loci reflect consistent ancestry-associated effects rather than population-specific signals and provides a stringent filter against false positives without requiring explicit correction. Significant windows pointing to the same genomic region across baselines were merged by overlap clustering into discrete loci.

### Candidate Gene Identification within Significant Loci

For each significant window, ancestry-associated genes were identified by refitting the logistic regression model at the gene level rather than the window level, restricting to genes whose genomic coordinates overlapped the significant window. Gene-level p-values were adjusted by Benjamini-Hochberg FDR correction within each baseline, and genes reaching q < 0.05 were retained as ancestry-associated genes. Genes are reported in order of ascending q-value within each baseline. Gene annotations were derived from GENCODE v43 (GRCh38).

### Ancestry-Directional ASE Analysis

To determine the direction of ancestry effects at significant loci (i.e. which ancestral haplotype is preferentially expressed), a complementary haplotype-resolved directional analysis was conducted. For each heterozygous site within a significant window, per-site log_2_ ancestry ratios were computed within each individual. Rather than labeling reads as reference or alternative allele (which is relative to the GRCh38 reference genome), reads were assigned to haplotypes based on local ancestry inference. This means that for an individual carrying one CEU haplotype and one YRI haplotype at a given locus, reads were grouped by which ancestral haplotype they originated from, and the log_2_ ratio of CEU-assigned reads to YRI-assigned reads was computed per site.

Sites were classified as anc1-dominant (log_2_ ratio > 0.5), anc2-dominant (log_2_ ratio < −0.5), or balanced (log_2_ ratio ≤ 0.5 and ≥ −0.5), with all classifications made per-site within each individual. At the window level (aggregated across individuals), an exact binomial test (null proportion 0.5) was applied to the number of anc1-dominant relative to anc2-dominant site observations pooled across all individuals in the window, and windows with a mean log2 ratio exceeding ±0.3 and a binomial p < 0.05 were classified as directionally biased. Results were restricted to windows with at least 10 individuals and at least 50 informative sites. Loci in which more than 70% or fewer than 30% of dominant calls carried the GRCh38 reference allele were flagged for potential reference allele mapping bias, as GRCh38 is a biased mosaic derived primarily from a single individual, with recognized limitations in representing global genetic diversity (Liao et al., 2023). To summarize direction at the 33 CEU/CEU scan loci (Table 3), we also calculated, for each locus, the percentage of informative genes with higher expression from the CEU haplotype (% CEU) and from the non-CEU haplotype (% YRI) among heterozygous individuals. A locus was labeled CEU-dominant when the CEU haplotype was higher for more than 60% of informative genes and balanced or mixed otherwise. The population-specific percentages reported in Results were calculated the same way within each population group, using loci with at least 20 informative genes.

## Results

### Allele-Specific Expression is Wide-Spread in a Multi-Ancestry Sample

The study design, data processing pipeline, and the three complementary analyses are summarized in Figure 1. We analyzed whole-blood RNA-sequencing data from 779 individuals (Table 1), spanning four major ascertainment-based groups: non-Hispanic White (NHW, n = 210), African American (AA, n = 193), Caribbean Hispanic (CHISP, n = 293), and Peruvian (PER, n = 83). Global ancestry estimates confirmed the expected ancestral diversity within each cohort, revealing significant intra-group variation in continental proportions (e.g., ranging from 39.9% to 97.2% YRI ancestry in the African American group, see Table 1).

**Figure 1.**
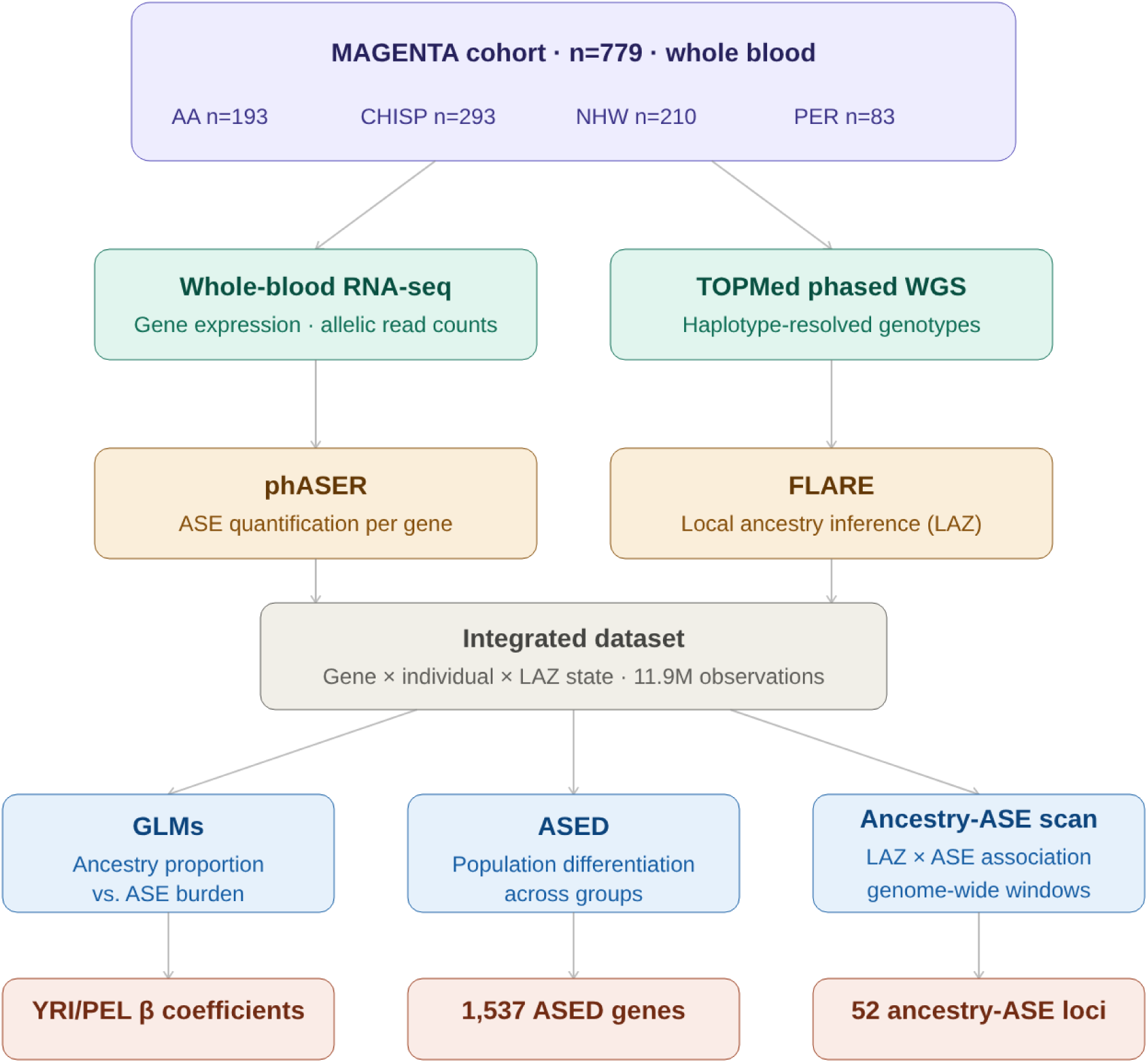
Study design and analytical pipeline. Whole-blood RNA-seq and TOPMed haplotype-resolved whole-genome sequencing data from 779 MAGENTA participants (African American n=193, Caribbean Hispanic n=293, Non-Hispanic White n=210, Peruvian n=83) were processed using phASER for allele-specific expression (ASE) quantification and FLARE for local ancestry inference, yielding 11.9 million gene-individual observations annotated with local ancestry zygosity (LAZ) states. Three complementary analyses were conducted on the integrated dataset: genome-wide generalized linear models (GLMs) relating ancestry proportion to ASE burden, a population differentiation test (ASED) identifying genes with inter-group ASE heterogeneity, and a genome-wide ancestry-ASE scan associating LAZ configurations with allelic imbalance across 1 Mb windows.

**Table 1.** Demographic and ASE summary characteristics of the MAGENTA study cohort.

| Characteristic | Population group |  |  |  | Overall<br>(N = 779) |
|---|---|---|---|---|---|
|  | Non-Hispanic White<br>(NHW, n = 210) | African American<br>(AA, n = 193) | Caribbean Hispanic<br>(CHISP, n = 293) | Peruvian<br>(PER, n = 83) |  |
| **DEMOGRAPHICS** |  |  |  |  |  |
| Age, median (range) | 78 (65–97) | 76 (63–97) | 77 (54–101) | 71 (53–92) | 77 (53–101) |
| Female, n (%) | 115 (54.8%) | 138 (71.5%) | 212 (72.4%) | 47 (56.6%) | 512 (65.7%) |
| AD, n (%) | 105 (50.0%) | 95 (49.2%) | 117 (39.9%) | 40 (48.2%) | 357 (45.8%) |
| Cognitively unimpaired, n (%) | 105 (50.0%) | 98 (50.8%) | 176 (60.1%) | 43 (51.8%) | 422 (54.2%) |
| **GLOBAL ANCESTRY** |  |  |  |  |  |
| YRI proportion, mean (SD) [range] | 0.013 (0.026)<br>[0.003–0.330] | 0.804 (0.124)<br>[0.399–0.972] | 0.182 (0.153)<br>[0.034–0.820] | 0.051 (0.057)<br>[0.006–0.302] | 0.276 (0.331)<br>[0.003–0.972] |
| PEL proportion, mean (SD) [range] | 0.007 (0.048)<br>[0.000–0.597] | 0.013 (0.010)<br>[0.002–0.058] | 0.165 (0.051)<br>[0.044–0.333] | 0.675 (0.188)<br>[0.091–0.912] | 0.139 (0.212)<br>[0.000–0.912] |
| **ASE BURDEN — ALL GENES** |  |  |  |  |  |
| Heterozygous genes, median (range) | 14,129 (4,900–21,359) | 16,974 (7,431–24,120) | 15,138 (2,479–22,477) | 13,922 (4,044–20,904) | 15,104 (2,479–24,120) |
| ASE-positive genes, median (range) | 5,901 (3,462–9,268) | 7,480 (5,424–10,072) | 7,187 (1,159–11,442) | 6,250 (1,914–10,935) | 6,848 (1,159–11,442) |
| **ASE BURDEN — PROTEIN-CODING GENES** |  |  |  |  |  |
| Heterozygous genes, median (range) | 9,907 (3,914–12,526) | 11,638 (5,836–14,326) | 10,553 (1,789–13,403) | 9,416 (2,969–12,632) | 10,465 (1,789–14,326) |
| ASE-positive genes, median (range) | 4,864 (2,970–6,654) | 6,144 (4,525–8,136) | 5,876 (978–8,932) | 5,126 (1,604–8,126) | 5,625 (978–8,932) |

AD = Alzheimer disease; ASE = allele-specific expression; YRI = Yoruba in Ibadan (African ancestry proxy); PEL = Peruvians in Lima (Amerindian ancestry proxy). Age reported as median (range). Cognitively unimpaired includes participants with no cognitive impairment diagnosis. Global ancestry proportions estimated by FLARE from TOPMed-imputed genotype data. ASE-positive genes were called as described in the Methods (at least 8 total reads per gene and individual, exact binomial test, FDR q ≤ 0.05). Heterozygous genes have at least one phased heterozygous site in that individual.

The sample was approximately 66% female overall, with group-specific proportions ranging from ∼55% (NHW) to ∼72% (AA and CHISP). Ages ranged from 53 to 101 years (rounded to whole numbers throughout), reflecting an older population relevant to late-onset traits. Median ages were similar across groups (NHW = 78, AA = 76, CHISP = 77, PER = 71 years). Alzheimer’s disease (AD) cases and cognitively normal individuals were included, with mild cognitive impairment (MCI) participants and 11 participants with other diagnoses collapsed into the cognitively unimpaired group given the intermediate and heterogeneous nature of MCI status. However, disease status was not the primary focus of our analyses. Across individuals, the median number of expressed genes containing at least one heterozygous variant (that is, genes with at least one phased heterozygous site in that individual) per sample ranged from 13,922 (PER) to 16,974 (AA), and the median number of ASE-positive genes ranged from 5,901 (NHW) to 7,480 (AA). Similar trends were observed when restricting to protein-coding genes. Differences by group likely reflect a combination of ancestry-related variation in allele frequency and sample-to-sample variability in gene expression (Table 1).

Using phased genotypes to assign RNA-seq reads to the two haplotypes, we quantified allelic imbalance for each gene in each individual. As expected, imbalance detection correlated with read depth and minor allele frequency. Of 11.9 million gene-individual observations across the cohort, 9.2 million (77%) had at least the 8 total reads required for ASE calling. Of these, 7.7 million had reads from both haplotypes and were tested with the binomial test, and 1.5 million had reads from a single haplotype. In total, 4.5 million observations (49% of those with at least 8 reads) were ASE-positive. Heterozygous gene count and ASE-positive gene count were strongly correlated across individuals (Spearman ρ = 0.63, p < 2.2×10⁻¹⁶), confirming that ASE detection sensitivity scales with the number of sites available for allele-specific measurement.

Within each individual, we quantified ASE burden by counting the number of ASE-positive genes. Prior work has shown that ASE detection sensitivity, read depth, mapping biases, and expression stability differ substantially between protein-coding versus non-protein-coding genes and between biallelic versus monoallelic ASE events (Castel et al., 2015), and that these features reflect both technical and biological sources of variability (Castel et al., 2020). Based on these considerations, we defined three outcome partitions across which we estimate ASE burden: all ASE-positive genes, ASE-positive protein-coding genes, and ASE-positive non-protein-coding genes. Of these, ASE-positive protein-coding gene count is treated as the primary outcome given its biological interpretability and well-characterized regulatory annotation, with each person having a median of 5,625 ASE-positive protein-coding genes (Table 1). Using this information, we then performed a set of analyses that examine how ASE burden differs by ancestry proportion and population group, followed by analyses that examine the influence of ancestry on a deeper haplotype-specific level.

### Genome-wide Analysis Reveals West African Ancestry Associates with Increased ASE Burden

Using the full sample of 779 individuals, we fit quasi-Poisson generalized linear models (GLMs) relating individual-level attributes to the ASE-positive gene count for each person. Individual-level local ancestry estimates provided continuous global ancestry proportions for European (CEU), West African (YRI), and Peruvian/Amerindian (PEL). We tested YRI and PEL ancestry proportions as predictors in separate GLMs, alongside age, sex, AD status, APOE genotype, and the number of heterozygous sites. Model specifications are given in Methods (Equations 1 and 2). Results for all ASE-positive genes and ASE-positive protein-coding genes are shown in Figure 2, and results for all three partitions are in Table 2. Results for heterozygous gene counts in the matching partitions are shown in Figure S1 and Supplementary Table S7.

**Figure 2.**
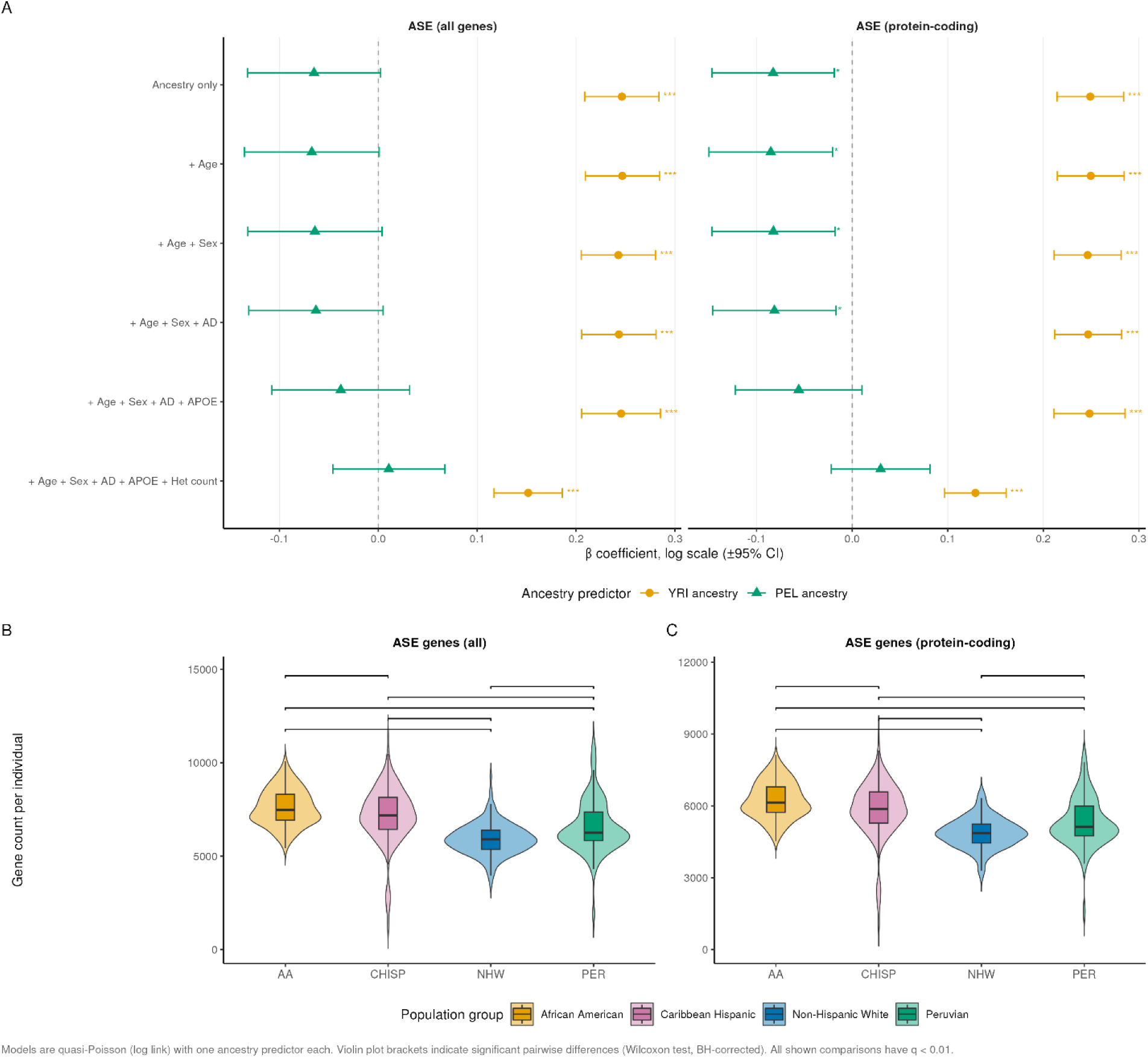
Ancestry predicts ASE-positive gene burden across population groups. (A) Beta coefficients (±95% CI) from generalized linear models predicting two ASE-positive gene count outcomes (all genes and protein-coding genes) as a function of YRI ancestry proportion (orange circles) or PEL ancestry proportion (green triangles), across six covariate adjustment sets, from ancestry alone to the fully adjusted model that also includes the number of heterozygous genes. Asterisks denote significance: *p<0.05, **p<0.01, ***p<0.001. (B, C) Distribution of per-individual ASE-positive gene counts by population group for all genes (B) and protein-coding genes (C). Brackets indicate significant pairwise differences by Wilcoxon rank-sum test with Benjamini-Hochberg correction. All shown comparisons have q<0.01. AA, African American; CHISP, Caribbean Hispanic; NHW, Non-Hispanic White; PER, Peruvian.

**Table 2.** Quasi-Poisson GLM results: association of global ancestry proportions with ASE-positive gene counts across sequential covariate models (N = 779)

| Response variable | Predictor | Covariates included | β | SE | z | p-value | Pseudo-R² | Sig. |
|---|---|---|---|---|---|---|---|---|
| **ASE-positive genes (all)** | **YRI proportion** | **None (unadjusted)** | **+0.2463** | **0.0191** | **12.901** | **1.21×10<sup>−34</sup>** | **0.1654** | *** |
|  |  | **Age** | **+0.2468** | **0.0191** | **12.900** | **1.24×10<sup>−34</sup>** | **0.1656** | *** |
|  |  | **Age, sex** | **+0.2428** | **0.0191** | **12.678** | **1.32×10<sup>−33</sup>** | **0.1711** | *** |
|  |  | **Age, sex, AD status** | **+0.2433** | **0.0192** | **12.694** | **1.12×10<sup>−33</sup>** | **0.1721** | *** |
|  |  | **Age, sex, AD status, APOE** | **+0.2456** | **0.0203** | **12.070** | **7.68×10<sup>−31</sup>** | **0.1732** | *** |
|  |  | **Age, sex, AD status, APOE, matched het gene count** | **+0.1516** | **0.0176** | **8.605** | **4.27×10<sup>−17</sup>** | **0.4374** | *** |
|  | PEL proportion | None (unadjusted) | -0.0650 | 0.0344 | -1.891 | 0.059 | 0.0044 |  |
|  |  | Age | -0.0672 | 0.0348 | -1.935 | 0.053 | 0.0046 |  |
|  |  | Age, sex | -0.0641 | 0.0346 | -1.853 | 0.064 | 0.0164 |  |
|  |  | Age, sex, AD status | -0.0631 | 0.0347 | -1.819 | 0.069 | 0.0168 |  |
|  |  | Age, sex, AD status, APOE | -0.0379 | 0.0356 | -1.067 | 0.286 | 0.0292 |  |
|  |  | Age, sex, AD status, APOE, matched het gene count | +0.0106 | 0.0288 | 0.368 | 0.713 | 0.3865 |  |
| **ASE-positive genes (protein-coding)** | **YRI proportion** | **None (unadjusted)** | **+0.2492** | **0.0178** | **13.992** | **7.74×10<sup>−40</sup>** | **0.1871** | *** |
|  |  | **Age** | **+0.2497** | **0.0178** | **13.994** | **7.65×10<sup>−40</sup>** | **0.1874** | *** |
|  |  | **Age, sex** | **+0.2464** | **0.0179** | **13.783** | **8.19×10<sup>−39</sup>** | **0.1915** | *** |
|  |  | **Age, sex, AD status** | **+0.2469** | **0.0179** | **13.802** | **6.68×10<sup>−39</sup>** | **0.1927** | *** |
|  |  | **Age, sex, AD status, APOE** | **+0.2483** | **0.0190** | **13.078** | **1.95×10<sup>−35</sup>** | **0.1940** | *** |
|  |  | **Age, sex, AD status, APOE, matched het gene count** | **+0.1290** | **0.0164** | **7.860** | **1.30×10<sup>−14</sup>** | **0.4830** | *** |
|  | **PEL proportion** | **None (unadjusted)** | **-0.0826** | **0.0326** | **-2.531** | **0.0116** | **0.0078** | * |
|  |  | **Age** | **-0.0851** | **0.0330** | **-2.577** | **0.0101** | **0.0081** | * |
|  |  | **Age, sex** | **-0.0824** | **0.0329** | **-2.504** | **0.0125** | **0.0182** | * |
|  |  | **Age, sex, AD status** | **-0.0813** | **0.0330** | **-2.468** | **0.0138** | **0.0186** | * |
|  |  | Age, sex, AD status, APOE | -0.0559 | 0.0338 | -1.655 | 0.098 | 0.0328 |  |
|  |  | Age, sex, AD status, APOE, matched het gene count | +0.0298 | 0.0263 | 1.133 | 0.258 | 0.4450 |  |
| **ASE-positive genes (non-protein-coding)** | **YRI proportion** | **None (unadjusted)** | **+0.2336** | **0.0266** | **8.797** | **8.97×10<sup>−18</sup>** | **0.0897** | *** |
|  |  | **Age** | **+0.2338** | **0.0266** | **8.788** | **9.70×10<sup>−18</sup>** | **0.0898** | *** |
|  |  | **Age, sex** | **+0.2266** | **0.0265** | **8.539** | **7.10×10<sup>−17</sup>** | **0.1003** | *** |
|  |  | **Age, sex, AD status** | **+0.2270** | **0.0266** | **8.547** | **6.70×10<sup>−17</sup>** | **0.1008** | *** |
|  |  | **Age, sex, AD status, APOE** | **+0.2330** | **0.0282** | **8.264** | **6.13×10<sup>−16</sup>** | **0.1016** | *** |
|  |  | **Age, sex, AD status, APOE, matched het gene count** | **+0.1619** | **0.0254** | **6.386** | **2.95×10<sup>−10</sup>** | **0.3062** | *** |
|  | PEL proportion | None (unadjusted) | +0.0128 | 0.0446 | 0.288 | 0.774 | 0.0001 |  |
|  |  | Age | +0.0114 | 0.0452 | 0.252 | 0.801 | 0.0002 |  |
|  |  | Age, sex | +0.0166 | 0.0448 | 0.370 | 0.712 | 0.0170 |  |
|  |  | Age, sex, AD status | +0.0177 | 0.0449 | 0.394 | 0.694 | 0.0172 |  |
|  |  | Age, sex, AD status, APOE | +0.0412 | 0.0463 | 0.891 | 0.373 | 0.0236 |  |
|  |  | Age, sex, AD status, APOE, matched het gene count | +0.0505 | 0.0406 | 1.243 | 0.214 | 0.2710 |  |

Significance codes: \* p < 0.05 \*\* p < 0.01 \*\*\* p < 0.001
β = quasi-Poisson regression coefficient (log scale), SE = standard error. Pseudo-R² = 1 − (residual deviance / null deviance) for the full model including all listed covariates plus the ancestry predictor. YRI = Yoruba in Ibadan (African ancestry proxy), PEL = Peruvians in Lima (Amerindian ancestry proxy). AD status was coded as AD versus cognitively unimpaired (MCI and other diagnoses included in the unimpaired group). The final covariate set for ASE outcomes adds the heterozygous gene count of the matching partition (all genes, protein-coding, or non-protein-coding). Each ancestry predictor was tested in a separate model. Bold text indicates p < 0.05. All models: N = 779.

**Table 3.** Summary of 33 ancestry-ASE scan loci identified under the CEU/CEU baseline (q < 0.05 in all four populations), ordered by effect size.

| Rank | Locus | Pattern | Best q-value | log<sub>2</sub>OR | % CEU | % YRI | Total genes | Protein-coding genes (n) | Protein-coding gene list |
|---|---|---|---|---|---|---|---|---|---|
| 1 | Chr7:140-141Mb | Balanced/Mixed | 0.0023 | 4.934 | 60.8% | 39.2% | 34 | 10 | TBXAS1, PARP12, KDM7A, SLC37A3, RAB19, MKRN1, DENND2A, ADCK2, NDUFB2, BRAF |
| 2 | Chr8:108-109Mb | Balanced/Mixed | 0.0012 | 4.179 | 46.3% | 53.7% | 14 | 4 | RSPO2, EIF3E, EMC2, TMEM74 |
| 3 | Chr20:31-32Mb | Balanced/Mixed | 9.05×10<sup>−11</sup> | 4.021 | 37.8% | 62.2% | 48 | 20 | DEFB115, DEFB116, DEFB118, DEFB119, DEFB121, DEFB123, DEFB124, REM1, HM13, MCTS2, ID1, COX4I2, BCL2L1, TPX2, MYLK2, FOXS1, DUSP15, TTLL9, PDRG1, XKR7 |
| **4** | **Chr12:110-111Mb** | **CEU-dominant** | **2.68×10<sup>−9</sup>** | **3.157** | **66.8%** | **33.2%** | **42** | **16** | **ANKRD13A, C12orf76, IFT81, ATP2A2, ANAPC7, ARPC3, GPN3, FAM216A, VPS29, RAD9B, PPTC7, TCTN1, HVCN1, PPP1CC, CCDC63, MYL2** |
| 5 | Chr13:45-46Mb | Balanced/Mixed | 7.41×10<sup>−5</sup> | 2.973 | 50.8% | 49.2% | 34 | 10 | GPALPP1, GTF2F2, KCTD4, TPT1, SLC25A30, COG3, ERICH6B, CBY2, SIAH3, ZC3H13 |
| 6 | Chr17:41-42Mb | Balanced/Mixed | 1.33×10<sup>−12</sup> | 2.488 | 50.8% | 49.2% | 98 | 61 | KRTAP3-1, KRTAP1-5, KRTAP1-4, KRTAP1-3, KRTAP1-1, KRTAP2-1, KRTAP2-2, KRTAP2-3, KRTAP2-4, KRTAP4-7, KRTAP4-8, KRTAP4-16, KRTAP4-9, KRTAP4-11, KRTAP4-12, KRTAP4-6, KRTAP4-5, KRTAP4-4, KRTAP4-3, KRTAP4-2, KRTAP4-1, KRTAP9-1, KRTAP9-2, KRTAP9-3, KRTAP9-8, KRTAP9-4, KRTAP9-9, KRTAP9-6, KRTAP9-7, KRTAP29-1, KRTAP16-1, KRTAP17-1, KRT33A, KRT33B, KRT34, KRT31, KRT37, KRT38, KRT32, KRT35, KRT36, KRT13, KRT15, KRT19, KRT9, KRT14, KRT16, KRT17, EIF1, GAST, HAP1, JUP, P3H4, FKBP10, NT5C3B, KLHL10, KLHL11, ACLY, ODAD4, CNP, DNAJC7 |
| 7 | Chr16:29-30Mb | Balanced/Mixed | 9.85×10<sup>−66</sup> | 2.383 | 57.9% | 42.1% | 90 | 25 | NPIPB11, BOLA2-SMG1P6, BOLA2, SLX1B, SULT1A4, NPIPB12, SPN, QPRT, C16orf54, ZG16, KIF22, MAZ, PRRT2, PAGR1, MVP, CDIPT, SEZ6L2, ASPHD1, KCTD13, TMEM219, TAOK2, HIRIP3, INO80E |
| 8 | Chr5:43-44Mb | Balanced/Mixed | 2.61×10<sup>−6</sup> | 1.888 | 42.4% | 57.6% | 31 | 9 | ANXA2R, ZNF131, NIM1K, HMGCS1, CCL28, TMEM267, C5orf34, PAIP1, NNT |
| 9 | Chr18:21-22Mb | Balanced/Mixed | 3.28×10<sup>−9</sup> | 1.725 | 62.1% | 37.9% | 34 | 7 | ROCK1, GREB1L, ESCO1, SNRPD1, ABHD3, MIB1 |
| 10 | Chr13:40-41Mb | Balanced/Mixed | 1.18×10<sup>−21</sup> | 1.715 | 43.7% | 56.3% | 31 | 4 | FOXO1, MRPS31, SLC25A15, ELF1 |
| 11 | Chr13:102-103Mb | Balanced/Mixed | 1.74×10<sup>−9</sup> | 1.707 | 52.6% | 47.4% | 27 | 10 | FGF14, TPP2, METTL21C, CCDC168, TEX30, POGLUT2, BIVM, BIVM-ERCC5, ERCC5, METTL21EP |
| 12 | Chr10:98-99Mb | Balanced/Mixed | 3.64×10<sup>−7</sup> | 1.678 | 51.2% | 48.8% | 16 | 7 | CRTAC1, R3HCC1L, LOXL4, PYROXD2, HPS1, HPSE2 |
| 13 | Chr15:51-52Mb | Balanced/Mixed | 1.91×10<sup>−17</sup> | 1.677 | 52.5% | 47.5% | 40 | 11 | AP4E1, TNFAIP8L3, CYP19A1, GLDN, DMXL2, SCG3, LYSMD2, TMOD2, TMOD3, LEO1, MAPK6 |
| 14 | Chr2:196-197Mb | Balanced/Mixed | 7.44×10<sup>−12</sup> | 1.528 | 38.7% | 61.3% | 19 | 8 | DNAH7, STK17B, HECW2, CCDC150, GTF3C3, C2orf66, PGAP1, ANKRD44 |
| 15 | Chr4:10-11Mb | Balanced/Mixed | 4.76×10<sup>−19</sup> | 1.505 | 47.7% | 52.3% | 29 | 4 | SLC2A9, WDR1, ZNF518B, CLNK |
| 16 | Chr17:20-21Mb | Balanced/Mixed | 2.72×10<sup>−6</sup> | 1.454 | 43.6% | 56.4% | 82 | 5 | SPECC1, LGALS9B, CDRT15L2, MEIS3P2, USP22 |
| 17 | Chr9:113-114Mb | Balanced/Mixed | 3.35×10<sup>−5</sup> | 1.339 | 61.1% | 38.9% | 33 | 16 | ZNF883, ZFP37, SLC31A2, FKBP15, SLC31A1, CDC26, PRPF4, RNF183, WDR31, BSPRY, HDHD3, ALAD, POLE3, C9orf43, RGS3, ZNF618 |
| 18 | Chr21:38-39Mb | Balanced/Mixed | 1.16×10<sup>−18</sup> | 1.313 | 39.9% | 60.1% | 35 | 4 | KCNJ6, KCNJ15, ERG, ETS2 |
| 19 | Chr8:27-28Mb | Balanced/Mixed | 1.44×10<sup>−5</sup> | 1.300 | 55.9% | 44.1% | 37 | 11 | STMN4, TRIM35, PTK2B, CHRNA2, EPHX2, CLU, SCARA3, CCDC25, ESCO2, PBK, SCARA5 |
| 20 | Chr3:120-121Mb | Balanced/Mixed | 0.0012 | 1.198 | 52.5% | 47.5% | 35 | 9 | GSK3B, GPR156, LRRC58, FSTL1, NDUFB4, HGD, RABL3, GTF2E1, STXBP5L |
| 21 | Chr16:53-54Mb | Balanced/Mixed | 2.70×10<sup>−9</sup> | 1.190 | 56.7% | 43.3% | 33 | 6 | CHD9NB, CHD9, RBL2, AKTIP, RPGRIP1L, FTO |
| 22 | Chr22:23-24Mb | Balanced/Mixed | 4.39×10<sup>−14</sup> | 1.176 | 55.6% | 44.4% | 76 | 22 | RSPH14, GNAZ, RAB36, BCR, IGLL1, DRICH1, RGL4, ZNF70, VPREB3, C22orf15, CHCHD10, MMP11, SMARCB1, DERL3, SLC2A11, MIF, GSTT2B, DDTL, DDT, GSTT2, GSTT4 |
| 23 | Chr16:85-86Mb | Balanced/Mixed | 9.96×10<sup>−8</sup> | 1.173 | 52.9% | 47.1% | 52 | 9 | ZDHHC7, KIAA0513, CIBAR2, GSE1, GINS2, C16orf74, EMC8, COX4I1, IRF8 |
| 24 | Chr6:31-32Mb | Balanced/Mixed | 1.19×10<sup>−28</sup> | 1.173 | 53.0% | 47.0% | 135 | 63 | MUC22, C6orf15, PSORS1C1, CDSN, PSORS1C2, CCHCR1, TCF19, POU5F1, HLA-C, HLA-B, MICA, MICB, MCCD1, DDX39B, ATP6V1G2, NFKBIL1, LTA, TNF, LTB, LST1, NCR3, AIF1, PRRC2A, BAG6, APOM, C6orf47, GPANK1, CSNK2B, LY6G5B, LY6G5C, ABHD16A, LY6G6F, LY6G6D, MPIG6B, LY6G6C, DDAH2, CLIC1, MSH5, SAPCD1, VWA7, VARS1, LSM2, HSPA1L, HSPA1A, HSPA1B, NEU1, SLC44A4, EHMT2, C2, ZBTB12, CFB, NELFE, SKIC2, DXO, WHR1, C4A [+7 more] |
| 25 | Chr19:39-40Mb | Balanced/Mixed | 3.21×10<sup>−6</sup> | 1.165 | 50.6% | 49.4% | 78 | 34 | FBXO27, ACP7, PAK4, NCCRP1, SYCN, IFNL3, IFNL4, IFNL2, IFNL1, LRFN1, GMFG, SAMD4B, PAF1, MED29, ZFP36, PLEKHG2, RPS16, SUPT5H, TIMM50, DLL3, SELENOV, EID2B, EID2, LGALS13, LGALS16, LGALS14, CLC, LEUTX, DYRK1B, FBL, FCGBP, PSMC4, ZNF546 [+1 more] |
| 26 | Chr3:23-24Mb | Balanced/Mixed | 1.19×10<sup>−4</sup> | 1.156 | 52.9% | 47.1% | 16 | 5 | UBE2E2, UBE2E1, NKIRAS1, RPL15, NR1D2 |
| 27 | Chr12:96-97Mb | Balanced/Mixed | 7.51×10<sup>−4</sup> | 1.137 | 47.5% | 52.5% | 24 | 5 | LTA4H, ELK3, CDK17, CFAP54, NEDD1 |
| 28 | Chr11:117-118Mb | Balanced/Mixed | 6.62×10<sup>−7</sup> | 1.129 | 41.9% | 58.1% | 35 | 14 | SIK3, PAFAH1B2, SIDT2, TAGLN, PCSK7, RNF214, BACE1, CEP164, DSCAML1, FXYD2, FXYD6, TMPRSS13, IL10RA [+1 more] |
| 29 | Chr22:19-20Mb | Balanced/Mixed | 1.10×10<sup>−11</sup> | 1.016 | 51.0% | 49.0% | 62 | 22 | FAM246C, DGCR2, ESS2, TSSK2, GSC2, SLC25A1, CLTCL1, HIRA, C22orf39, MRPL40, UFD1, CDC45, CLDN5, SEPTIN5, GP1BB, TBX1, GNB1L, RTL10, TXNRD2, COMT, ARVCF [+1 more] |
| 30 | Chr17:9-10Mb | Balanced/Mixed | 1.76×10<sup>−4</sup> | 0.893 | 49.8% | 50.2% | 25 | 9 | NTN1, STX8, CFAP52, USP43, DHRS7C, GSG1L2, GLP2R, RCVRN, GAS7 |
| 31 | Chr1:46-47Mb | Balanced/Mixed | 2.21×10<sup>−7</sup> | 0.820 | 53.5% | 46.5% | 44 | 22 | MAST2, PIK3R3, TSPAN1, POMGNT1, LURAP1, RAD54L, LRRC41, UQCRH, NSUN4, FAAH, DMBX1, TMEM275, KNCN, MKNK1, MOB3C, ATPAF1, TEX38, EFCAB14, CYP4B1, CYP4A11 [+2 more] |
| 32 | Chr15:63-64Mb | Balanced/Mixed | 7.10×10<sup>−12</sup> | 0.678 | 56.1% | 43.9% | 30 | 10 | TPM1, LACTB, RPS27L, RAB8B, APH1B, CA12, USP3, FBXL22, HERC1, DAPK2 |
| 33 | Chr16:0-1Mb | Balanced/Mixed | 2.63×10<sup>−32</sup> | 0.516 | 56.1% | 43.9% | 115 | 48 | POLR3K, SNRNP25, RHBDF1, MPG, NPRL3, HBZ, HBM, HBA2, HBA1, HBQ1, LUC7L, FAM234A, RGS11, ARHGDIG, PDIA2, AXIN1, MRPL28, PGAP6, NME4, DECR2, RAB11FIP3, CAPN15, PRR35, PIGQ, NHLRC4, RAB40C, WFIKKN1, METTL26, MCRIP2, WDR90, RHOT2, RHBDL1, STUB1, JMJD8, WDR24, FBXL16, METRN, ANTKMT, CCDC78, HAGHL, CIAO3, MSLN, MSLNL, RPUSD1, CHTF18, GNG13, LMF1 [+1 more] |

Loci are ordered by effect size (log<sub>2</sub>OR) in descending order. Pattern: Balanced/Mixed = neither ancestral haplotype consistently dominant across heterozygous individuals; CEU-dominant = CEU haplotype shows higher ASE rate in >60% of heterozygous individuals (blue shading). % CEU and % YRI reflect the mean percentage of heterozygous individuals at the locus showing higher expression from the CEU or YRI haplotype respectively. Best q-value = FDR-adjusted p-value from the most significant 1 Mb window within the locus under the CEU/CEU baseline. log<sub>2</sub>OR = log<sub>2</sub> odds ratio from the logistic regression at the best window. Total genes = all genes overlapping the locus window(s). Protein-coding gene list reflects genes present in all four population groups. All 33 loci were significant (q < 0.05) under the CEU/CEU baseline in all four populations.

YRI ancestry proportion showed a consistently positive and statistically significant association with ASE-positive gene counts (Figure 2). The effect was robust across all covariate adjustments and was strongest for protein-coding gene subsets, which exhibited the highest pseudo-R² values (0.187 to 0.194 for ASE-positive protein-coding gene counts), likely reflecting the greater read depth and detection sensitivity for protein-coding genes (Castel et al., 2015). Notably, YRI ancestry proportion remained a significant predictor of ASE-positive protein-coding gene count even after adjusting for the number of heterozygous protein-coding genes (β = 0.129, p = 1.3×10⁻¹⁴, R² = 0.483, compared with 0.444 for the same model without YRI ancestry, Supplementary Table S2), confirming that the ancestry effect on ASE burden is not attributable solely to admixed individuals carrying more testable sites due to increased genetic diversity in African-ancestry populations.

In the model comparison described in Methods, YRI ancestry proportion alone explained substantially more variance in ASE-positive protein-coding gene counts than all other covariates combined (R² = 0.187 versus R² = 0.030), with the fully adjusted model explaining only marginally more variance than ancestry alone (R² = 0.194). The association of YRI ancestry with ASE-positive gene counts remained significant upon inclusion of age, sex, AD status, and APOE genotype, with the effect size (β) for YRI ancestry shifting by less than 5% across all covariate models (range of log-scale coefficients: 0.246 to 0.250 for protein-coding ASE genes), indicating that the ancestry effect is not confounded by these clinical and demographic variables.

PEL ancestry proportion showed no significant association with ASE-positive gene counts across any partition in the fully adjusted model (p = 0.098 to 0.373), although the protein-coding association reached nominal significance (p < 0.05) before adjustment for APOE genotype and was associated with reduced heterozygous gene counts for all genes and protein-coding genes (β = −0.104 for protein-coding heterozygous count, p = 5.0×10^-^⁴, Supplementary Table S7). This pattern is consistent with reports from 1000 Genomes showing lower average number of variant sites in PEL genomes relative to other populations (Auton et al., 2015). Together, these results indicate that ancestry-linked differences in ASE burden (Price et al., 2008; Reich et al., 2012) are driven primarily by the YRI component of ancestry, which captures a major axis of genomic diversity known to influence allele frequencies, eQTL architecture, and expression variance (Stranger et al., 2007, Aguet et al., 2017).

Other covariates including age, sex, AD status, and APOE genotype had minimal impact on any outcome and did not alter the magnitude or direction of ancestry effects. Prior studies have reported modest associations between age and transcriptional instability (Bahar et al., 2006) or sex-specific regulatory variation (Aguet et al., 2020). In our data, age showed no significant association with ASE burden for all genes and protein-coding genes (p = 0.43 and 0.35 respectively in the fully adjusted model). Sex showed a modest negative association with ASE-positive gene counts (β = −0.032, p = 0.023 for all genes and β = −0.026, p = 0.045 for protein-coding genes, on the log scale), with male individuals carrying approximately 3% fewer ASE-positive genes on average. AD status and *APOE* genotype, while relevant to disease-specific expression differences (Mathys et al., 2019), did not meaningfully explain variance in ASE burden once ancestry proportions were accounted for. Comparisons of linear versus quadratic ancestry models revealed no evidence of non-linearity, indicating that a linear framework was sufficient to capture ancestry-linked variation in ASE count burdens. Full model estimates, confidence intervals, and statistical summaries for the ancestry terms are presented in Table 2, and coefficients for all covariates are provided in Supplementary Table S4.

We next examined ASE burden across populations to identify specific genes that may account for differences in ASE burden by ancestry. We estimated pairwise group-level differences in ASE by applying a gene-level adaptation of the ASED framework (Tian et al., 2018). For each gene, we computed the per-group allelic imbalance as the absolute difference between allele-specific counts divided by total reads at heterozygous positions. Among 15,008 genes tested, 2,053 showed significant between-group differences in allelic imbalance at 5% FDR, and 1,537 of these had at least 20 informative individuals in each of the four groups and are reported in Supplementary Table S1. Pairwise Wilcoxon tests indicated the strongest contrast was between AA vs NHW, followed by CHISP vs PER. In the AA vs NHW contrast, African American individuals showed greater median allelic imbalance, consistent with the higher YRI ancestry proportion in that group. Notably, the major histocompatibility complex (MHC) emerges across all four population groups (Supplementary Table S1) (**Figure 3**), consistent with the extensive population-specific regulatory variation in this region (Lam et al., 2017; Gutierrez-Arcelus et al., 2020).

**Figure 3.**
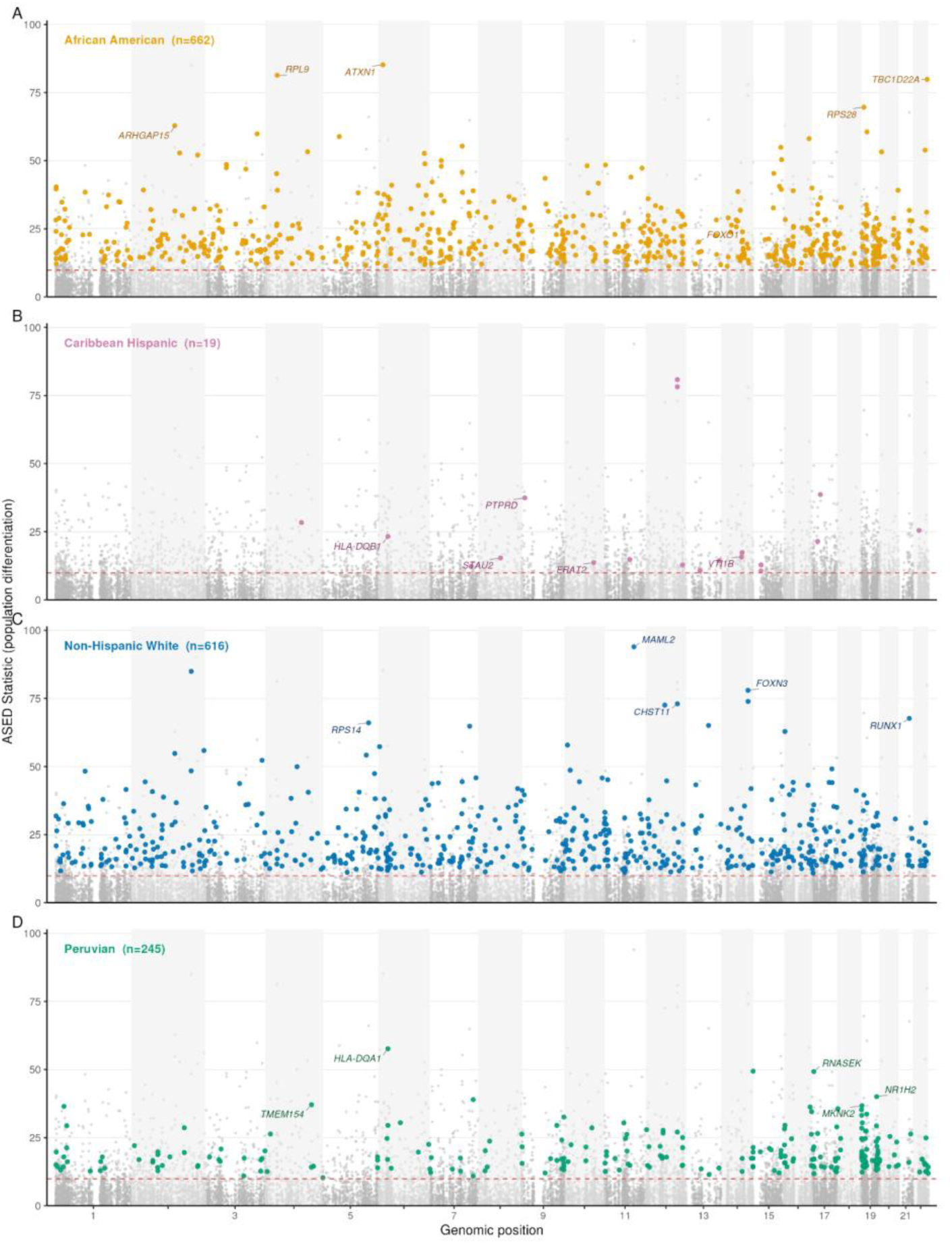
Genome-wide ASED Manhattan plot by population with the highest imbalance. Each point represents one gene tested for inter-group ASE heterogeneity, plotted by genomic position (x-axis) and ASED statistic (y-axis), a measure of population differentiation in allele-specific expression analogous to FST. Points are colored by the population group with the highest median allelic imbalance for that gene. Genes tied between groups appear in each tied group. Grey points indicate non-significant genes (q≥0.05 by Benjamini-Hochberg correction). The dashed red line marks the 5% FDR threshold. Only protein-coding genes with assigned gene symbols are labeled. n=1,537 significant genes across all four panels.

### Window-based Scans Identify Regions and Genes with Increased ASE Events by Ancestry

Motivated by the possibility of imprinting control regions or similar mechanisms that may impart ASE effects over a collection of genes, we next sought to identify genome regions that are enriched for ancestry-associated ASE events. We first characterized the local ancestry zygosity (LAZ) across the genome in 1 Mb non-overlapping windows. We then performed logistic regression analyses to model the presence of one or more ASE events for each LAZ-heterozygous configuration relative to each of three reference baselines (CEU/CEU, YRI/YRI, and PEL/PEL). The analysis proceeds in three steps: a genome-wide scan of windows to identify ancestry-associated loci, a gene-level refit within each significant locus to identify the ancestry-associated genes (Figure 6), and a directional analysis to determine which ancestral haplotype is more highly expressed (Figure S2).

We tested 2,213 1-Mb windows across the genome with sufficient local ancestry variation (with a minimum of 200 variant-level observations per window). The scan identified 656, 461, and 564 windows with FDR-significant ancestry effects (q < 0.05) in at least two of the four population groups under the CEU/CEU, YRI/YRI, and PEL/PEL baselines, respectively. Windows were further required to reach significance in all four population groups within the same baseline, ensuring that identified loci reflect consistent ancestry-associated effects rather than population-specific signals. This filter reduced the window sets to 33, 11, and 16 replicated windows for the CEU/CEU, YRI/YRI, and PEL/PEL baselines, respectively (Figure 4, Table 3). Adjacent significant windows were merged, yielding a final set of 52 non-redundant ancestry-associated ASE loci (from 53 unique replicated windows) across all three baselines. This scan identified loci with known population-differentiated regulatory variation. Locus-level plots for selected regions (windows in the top row and genes in the bottom row) are shown in Figure 5. The 16p11.2 region reached q = 9.8×10^−66^ with individuals carrying a non-European haplotype showing higher rates of ASE than CEU/CEU homozygous individuals in this region. This region is a known hotspot for neurodevelopmental copy number variants (Golzio et al., 2012). The major histocompatibility complex (MHC, chr6:31–32 Mb), a region with well-characterized population-differentiated haplotype architecture and extensive regulatory variation (Lam et al., 2017; Kamitaki et al., 2020), was detected at q = 1.2 × 10^−28^. The beta-defensin cluster (chr20:31–32 Mb, labeled HM13 in Figure 4), which exhibits extensive copy number variation across human populations (Hollox et al., 2003; Hardwick et al., 2011), was also significant at q = 9.1 × 10^−11^ (CEU/CEU). Chr15:63–64 Mb and chr13:102–103 Mb were detected under both CEU/CEU and YRI/YRI baselines but not the PEL/PEL baseline, indicating replication across African and European reference configurations but not Amerindian ancestry.

**Figure 4.**
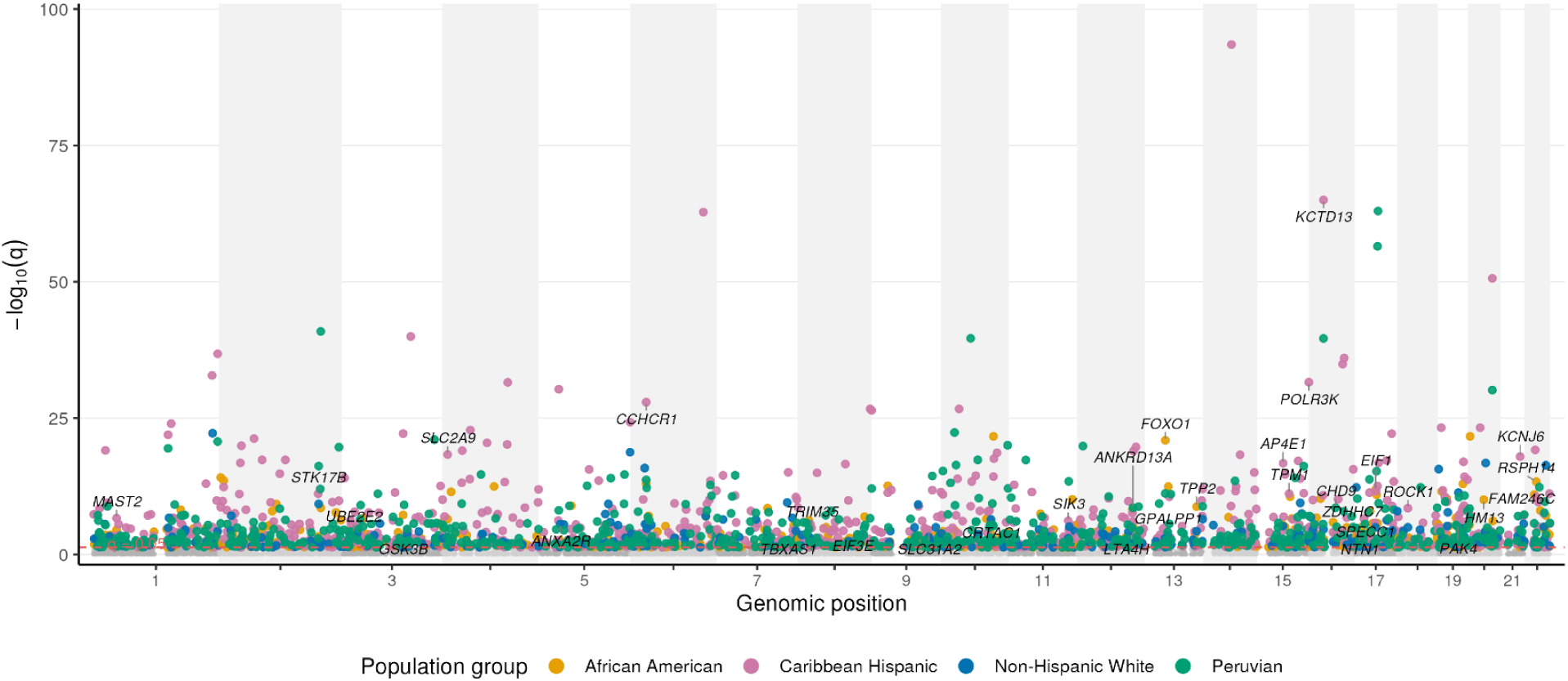
Genome-wide ancestry-ASE scan Manhattan plot by population group. Each point represents one 1 Mb genomic window tested in one population group for association between LAZ-heterozygous configuration and elevated ASE rates under the CEU/CEU baseline, plotted by genomic position (x-axis) and −log10(q) (y-axis). Points are colored by population group. Grey points indicate non-significant windows (q≥0.05). The dashed red line marks the 5% FDR threshold. Gene labels indicate the most significant protein-coding gene within each significant locus. Significant windows in adjacent positions were merged into 33 non-redundant loci that were significant in all four populations. AA, African American; CHISP, Caribbean Hispanic; NHW, Non-Hispanic White; PER, Peruvian.

**Figure 5.**
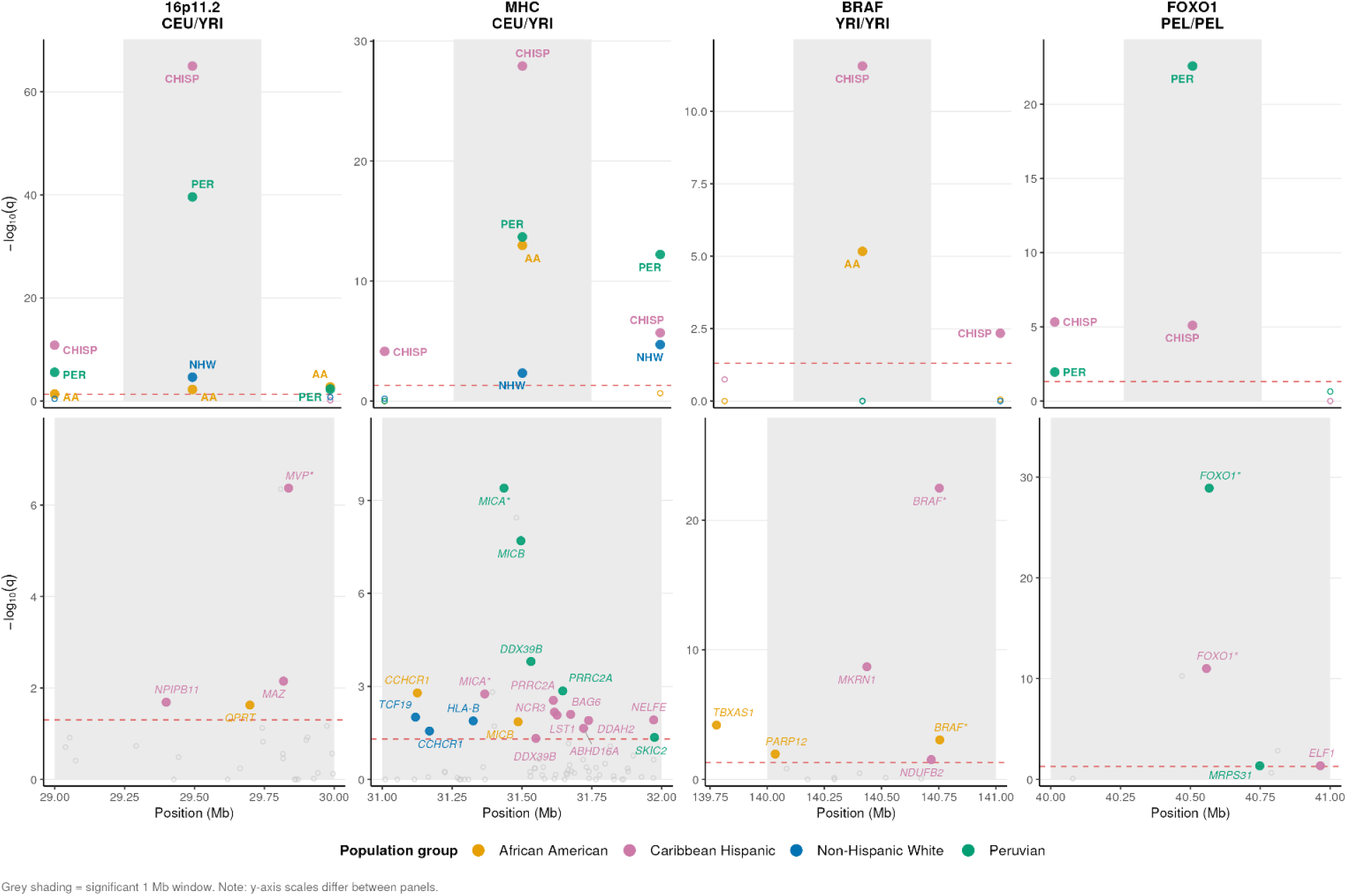
Locus zoom plots for four representative ancestry-ASE loci. Each column shows one significant locus. Top row: window-level scan signal (−log10(q)) across the locus. Grey shading marks the significant 1 Mb window. Each panel title gives the locus and the ancestry-pair term tested against the CEU/CEU baseline. Points are colored by population group, and population abbreviations label the significant windows for each group. Bottom row: gene-level association results within the significant window. Colored points indicate significant protein-coding ancestry-associated genes (q<0.05); grey open circles indicate all other tested genes. The asterisk (*) marks the protein-coding gene with the lowest q-value at the locus, and it is shown for each population in which that gene is significant. Note: y-axis scales differ between panels. Loci shown: 16p11.2 (Chr16:29–30 Mb), MHC (Chr6:31–32 Mb), BRAF region (Chr7:140–141 Mb), and FOXO1 region (Chr13:40– 41 Mb).

**Figure 6.**
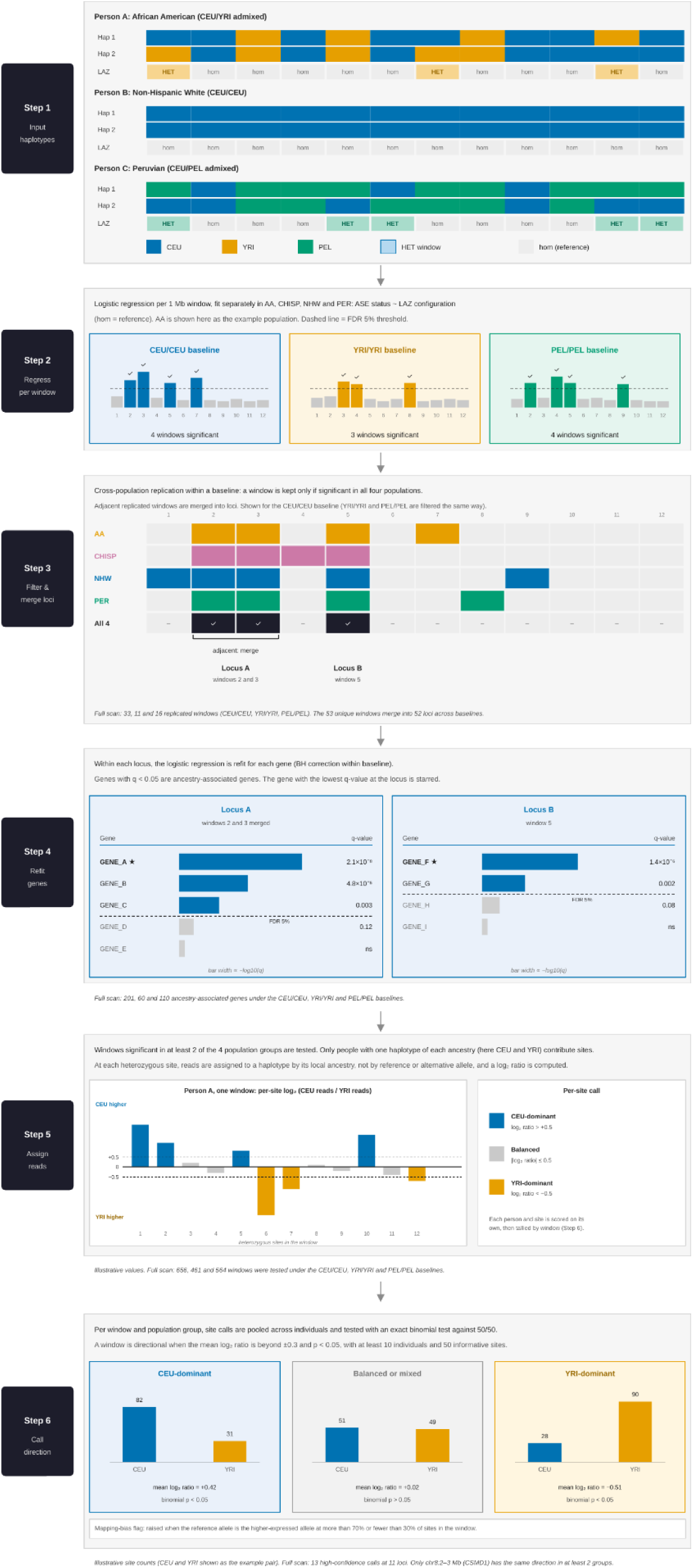
Schematic overview of the ancestry-ASE scan pipeline, from phased haplotype input to ancestry-associated gene identification and ancestry direction. (A) Input: phased haplotypes and local ancestry calls. For each individual, phased genotype data are integrated with local ancestry inference (FLARE) to assign an ancestral origin (CEU for non-Hispanic White-associated, YRI for African-associated, or PEL for Peruvian-associated) to each haplotype across non-overlapping 1 Mb genomic windows. Three representative individuals are shown: an African American individual with CEU/YRI admixture (Person A), a non-Hispanic White individual homozygous for CEU ancestry across all windows (Person B), and a Peruvian individual with CEU/PEL admixture (Person C). Each window is classified by its local ancestry zygosity (LAZ): heterozygous (HET, two different ancestral haplotypes) or homozygous (hom, both haplotypes from the same ancestry). Only HET windows contribute informative observations to the scan; hom windows serve as the reference category. **(B) Regression per window.** For each non-overlapping 1 Mb window and each ancestry-pair baseline (CEU/CEU, YRI/YRI, and PEL/PEL), a logistic regression, fit separately in each population group (AA, CHISP, NHW, and PER), tests whether individuals heterozygous for that ancestry pair (LAZ-HET) show higher rates of allele-specific expression than homozygous individuals (LAZ-hom, reference). The y-axis shows –log₁₀(q-value), and the dashed line marks the 5% FDR significance threshold. Colored bars indicate windows reaching significance (✓) under each baseline and gray bars are non-significant. AA is shown as the example population. Each baseline scan is performed independently across all windows genome-wide. **(C) Cross-population replication filter and locus merging.** Within each baseline, a window is retained only if it reaches significance (q < 0.05) in all four population groups, so that signals replicate across populations and are unlikely to reflect population-specific effects. The example shows the CEU/CEU baseline, in which windows 2, 3, and 5 are significant in all four populations. Retained windows that are adjacent are merged into loci, here windows 2 and 3 into Locus A, with window 5 as Locus B. In the full scan, this filter retained 33, 11, and 16 windows for the CEU/CEU, YRI/YRI, and PEL/PEL baselines. The 53 unique replicated windows merged into 52 non-redundant loci across the three baselines. **(D) Ancestry-associated gene identification within loci.** Within each locus, the logistic regression is refit for each gene whose coordinates overlap the replicated window, and p-values are adjusted by the Benjamini-Hochberg procedure within each baseline. Genes with q < 0.05 are ancestry-associated genes, and the gene with the lowest q-value at each locus is marked with a star (★). The horizontal bar length is proportional to –log₁₀(q), and the dashed line marks the 5% FDR threshold. Gene names shown are placeholders. In the full scan, 201, 60, and 110 ancestry-associated genes were identified under the CEU/CEU, YRI/YRI, and PEL/PEL baselines. **(E) Haplotype-resolved read assignment.** Windows significant in at least two of the four population groups (656, 461, and 564 windows under the CEU/CEU, YRI/YRI, and PEL/PEL baselines) are tested for direction. At each heterozygous site in an individual carrying one haplotype of each ancestry, reads are assigned to a haplotype by local ancestry rather than by reference or alternative allele, and a log₂ ratio of reads from the two ancestries is computed. Sites with a log₂ ratio above +0.5 are called dominant for the first ancestry (here CEU), sites below −0.5 are called dominant for the second (here YRI), and sites between these values are called balanced. The site values shown are illustrative. **(F) Ancestry direction.** Within each window and population group, dominant-site counts are pooled across individuals and compared with an equal split by an exact binomial test. A window is called directional when the mean log₂ ratio is beyond ±0.3 and the binomial p-value is below 0.05, and only windows with at least 10 individuals and 50 informative sites are evaluated. A window is flagged for possible mapping bias when the reference allele is the higher-expressed allele at more than 70% or fewer than 30% of sites. The three example windows show CEU-dominant, balanced, and YRI-dominant outcomes, and the counts shown are illustrative. In the full scan, 13 directional calls were made at 11 loci (Supplementary Table S5 and Supplementary Figure S2), and only chr8:2–3 Mb (CSMD1) showed the same direction in at least two population groups.

Each significant window spans a median of 35 genes, so to identify the specific genes responsible, we refit the logistic regression at gene resolution within each significant window. After Benjamini-Hochberg correction, this procedure identified 201, 60, and 110 ancestry-associated genes under the CEU/CEU, YRI/YRI, and PEL/PEL baselines, respectively, from 713, 177, and 318 significant gene-level tests (Supplementary Table S3). A gene can contribute more than one test when it is significant for different ancestry pairs or in different population groups. Most significant loci contained several ancestry-associated genes. Under the CEU/CEU baseline, the 33 loci contained a median of 4 ancestry-associated genes (range: 1–26), and three loci contained a single gene (EIF3E at chr8:108–109 Mb, TPP2 at chr13:102–103 Mb, and SPECC1 at chr17:20–21 Mb). Under YRI/YRI, the 11 loci contained a median of 6 genes (range: 1–12), with single-gene loci at chr2:26–27 Mb (RAB10) and chr13:102–103 Mb (TPP2). Under PEL/PEL, the 16 significant windows (15 merged loci) contained a median of 6.5 genes (range: 2– 17). Ancestry-associated ASE at most loci therefore involved a cluster of genes rather than a single focal gene. The locus with the most ancestry-associated genes (26 genes at chr6:31–32 Mb, from 94 significant tests) overlaps the gene-dense proximal MHC region. Full gene lists and all significant tests are provided in Supplementary Table S3. Among the most significant ancestry-associated genes compared to the CEU/CEU baseline was an unannotated lncRNA (ENSG00000291188) at 16p11.2 (q = 2.39 × 10^−50^). Among the top ancestry-associated genes were *FOXO1* under CEU/CEU (q = 1.22 × 10⁻²⁹), *USP3* under YRI/YRI (q = 8.14 × 10⁻⁴³), and *BRAF* under PEL/PEL (q = 3.79 × 10⁻¹⁸). *BRAF* was also significant under the CEU/CEU baseline (q = 3.28 × 10⁻²³), consistent with cross-baseline replication of the *BRAF* locus.

Using the CEU/CEU baseline as the primary reference, the 33 significant loci contained a median of 35 total genes per locus (range: 14–135) and a median of 10 protein-coding genes per locus (range: 4–63), and 30 of the 33 loci contained more than one ancestry-associated gene. These gene-level associations show which genes are linked to ancestry-associated ASE, but they do not show which ancestral haplotype is expressed more highly when ASE occurs. To address this, we next summarized the direction of expression at each locus. Of 33 loci, 32 (97%) showed a balanced or mixed directional pattern, indicating that neither the CEU nor the non-CEU haplotype consistently dominates expression across individuals at these regions. A single locus (Chr12:110– 111 Mb) showed a CEU-dominant pattern (66.8% CEU-biased), with 16 protein-coding genes including *MYL2*, *ATP2A2*, and *PPP1CC*. The window-level binomial test described below classified this window as balanced in every population group and baseline, so the CEU lean at this locus is modest. We also asked whether the loci with the strongest ancestry-ASE association were the ones where one haplotype was expressed more highly. They were not. The scan tests how often ASE occurs in people who carry two different ancestries at a window, whereas the directional analysis asks which of the two haplotypes contributes more of the expression. At the 33 CEU/CEU loci, no window with at least 10 individuals and 50 informative sites received a directional call in any population group, and the mean log₂ ratio of CEU to YRI reads stayed near zero (mean 0.008, range −0.39 to 0.23 across the 132 locus and population combinations). The average log₂ ratio at a locus was also unrelated to its scan effect size (Spearman ρ = −0.14, p = 0.43). Increased ASE at these loci therefore does not come with consistent preferential expression of one ancestral haplotype. The involvement of several genes at most loci and the predominantly balanced directional pattern suggest that ancestry-associated ASE at these regions often reflects distributed cis-regulatory architecture rather than focal variant effects (Wheeler et al., 2016).

Among African American individuals, where both CEU and YRI haplotypes are present at sufficient frequency to assess direction, the directional split between YRI-dominant and CEU-dominant genes was approximately equal (median 50.7% CEU-dominant, range 20.2 to 70.5% across loci with at least 20 informative genes). Among Caribbean Hispanic individuals the split was similarly balanced (median 51.0% CEU-dominant, range 23.3 to 77.9%). Among Peruvian individuals CEU and non-CEU haplotypes were nearly evenly split (median 52.6% CEU-dominant, range 8.3 to 91.9%), though the wider range reflects the smaller Peruvian sample size (n=83) and consequent variability in directional estimates at individual loci. These directional patterns were not genome-wide: mean log₂ ancestry ratios were near zero across all three baselines (range: 0.006 to 0.011), confirming that no single ancestral haplotype systematically dominates the admixed transcriptome.

### Preferential Expression of an Ancestral Haplotype at Some Genes

To determine the direction of ancestry effects at significant loci, we performed a complementary haplotype-resolved directional analysis, providing information on which ancestral haplotype is preferentially expressed. For each heterozygous site within a significant window, we computed per-site log_2_ ancestry ratios using read counts assigned by local ancestry rather than by reference/alternative allele identity (which is reference-genome-dependent and might limit ancestry assignment with GRCh38 reference bias). Windows were classified as ancestry1-dominant, ancestry2-dominant, or balanced using a combination of mean log_2_ ratio threshold (±0.3) and a binomial test for directional consistency across sites (p < 0.05), with reliability filters requiring ≥10 individuals and ≥50 informative sites per window.

At the global level, mean log_2_ ancestry ratios were indistinguishable from zero across all three baselines and all admixed populations (range: 0.000–0.024), indicating that neither ancestral cis-regulatory architecture systematically dominates in the admixed regulatory environment. This symmetry confirms that locus-specific directional signals reflect genuine regulatory disagreement rather than a genome-wide bias. Population-differentiated regulatory effects are therefore locus-specific phenomena: genome-wide, neither ancestry systematically dominates, but at specific loci, one haplotype architecture can be consistently preferential.

The directional analysis was applied to all windows significant in at least two of the four population groups (656, 461, and 564 windows under the CEU/CEU, YRI/YRI, and PEL/PEL baselines, respectively), rather than only the 33, 11, and 16 windows significant in all four. Among loci with high-confidence directional classification (n_samples ≥10, n_sites ≥50, direction ≠ balanced_or_mixed), 13 locus–direction calls at 11 unique loci showed consistent ancestry-specific dominance in at least one admixed population (Supplementary Table S5 and Figure S2). None of these loci overlapped the 53 replicated windows from any baseline (Table 3 and Supplementary Table S3), so the gene-level results do not apply to them. These loci are windows that were significant in at least two population groups but not in all four, so they were not replicated windows and were not carried into the gene-level refit. Applying a stricter cross-population replication criterion requiring the same locus to show the same direction independently in at least two populations, only one locus met this threshold: chr8:2–3 Mb (*CSMD1*), which showed PEL-dominant expression in both CHISP and PER under the PEL/PEL baseline. The same locus showed YRI-dominant expression in AA under the CEU/CEU and YRI/YRI baselines. The full directional classification results, including per-population direction calls, sample counts, and mapping bias flags, are provided in Supplementary Table S6. As a positive methodological control, chr15:48–49 Mb, containing *SLC24A5* which encodes a cation exchanger with one of the largest known ancestry-frequency differences between European and non-European populations (Lamason et al., 2005; Quillen et al., 2008) showed strong CEU-dominant expression (bias flag absent), validating the haplotype-resolved directional approach.

Allele frequency differences between populations do not automatically imply ASE. The 17q21 inversion locus (chr17:46–47 Mb), encompassing *MAPT*, *KANSL1*, *NSF*, and *WNT3*, showed CEU-dominant expression consistent with known H1/H2 expression differences across ancestry groups, though direct allele-specific expression at this locus has not been formally demonstrated (Donnelly et al., 2010; Pedicone et al., 2024). This call was specific to one population group under the CEU/CEU baseline. Caribbean Hispanic individuals carrying a CEU/YRI configuration at this window (n = 62, about 21% of that group) showed CEU-dominant expression (mean log₂ ratio = 0.36, bias flag absent). African Americans (n = 63, about 33% of that group) leaned the same way but fell just below the 0.3 threshold (mean log₂ ratio = 0.29). Non-Hispanic White and Peruvian individuals had too few carriers (n = 7 and n = 3) to assess direction, and the locus was balanced in every group under the PEL/PEL baseline. Because n counts individuals who contributed informative sites, these percentages are lower bounds on how common the configuration is. The MHC region (chr6:32–34 Mb) showed YRI-dominant and CEU-dominant calls depending on window and baseline, and all of these calls carry a mapping bias flag, so its direction should be interpreted with caution given the extensive haplotype-specific diversity at this locus (Gutierrez-Arcelus et al., 2020).

The most notable finding from the directional analysis was the *CSMD1* locus (chr8:2–3 Mb). *CSMD1*, one of the largest genes in the human genome, encodes CUB and Sushi multiple domains 1 with high expression in brain tissue and genetic associations with schizophrenia and cognitive traits (Escudero-Esparza et al., 2013; Havik et al., 2011). Under the CEU/CEU baseline, YRI-ancestry haplotypes were preferentially expressed in AA (n = 92, mean log_2_ ratio = −1.03, q = 1.41 × 10^−25^, bias flag absent). A consistent pattern was observed when using the PEL/PEL baseline rather than the CEU/CEU baseline: Under the PEL/PEL baseline, PEL-ancestry haplotypes were preferentially expressed in CHISP (n = 117, mean log_2_ ratio = −0.76, q = 2.25 × 10^−40^, bias flag absent) and PER (n = 42, mean log_2_ ratio = −1.67, q = 8.92 × 10^−99^, bias flag absent). Across both baselines, the European-derived haplotype was consistently not the dominant haplotype at this locus regardless of which non-European ancestry was present, pointing to a locus-specific regulatory pattern in which the European-derived haplotype is relatively less expressed when paired with either African or Amerindian haplotypes. This finding does not imply a rank ordering among non-European ancestries, as YRI and PEL haplotypes were never directly compared at this locus.

Importantly, the *CSMD1* directional signal was not captured by the gene-level association analysis, demonstrating that ancestry-specific regulatory dominance represents a dimension of ancestry-associated regulation independent of overall admixture-associated ASE burden. Directional bias analysis thus identifies loci not identified by the gene-level analysis, providing complementary and non-redundant biological information. The mechanism underlying CEU haplotype suppression at *CSMD1* is not yet established, but candidate explanations include population-differentiated cis-regulatory variants in the promoter or enhancer elements of *CSMD1* that reduce transcriptional activity on the European-derived haplotype, or differences in chromatin accessibility between ancestral backgrounds that alter transcription factor occupancy. The large size of CSMD1 (spanning over 2 Mb) and its high expression in neural tissue suggest that the detected regulatory effect in whole blood may reflect a subset of the regulatory complexity at this locus. Functional follow-up in tissue-specific or single-cell contexts would be required to determine the mechanistic basis.

## Discussion

This study integrates allele-specific expression with locus-resolved local ancestry inference to characterize the regulatory consequences of admixture in a multi-ancestry cohort of older adults. Across a series of complementary analyses spanning global ASE burden, gene-level population differences, and a genome-wide scan explicitly designed around ancestry configuration, we find that genetic ancestry shapes gene regulation at multiple scales. Collectively, these results establish that local ancestry is not merely a confounder to be adjusted away in genomics studies, but an informative predictor of allelic regulatory state that can reveal functional divergence invisible to standard population-level approaches.

At the broadest scale, inter-individual variation in ASE burden is driven primarily by the YRI component of global ancestry, with a consistently positive association across all tested gene categories. This effect was robust across covariate models and was not substantially modified by age, sex, Alzheimer’s disease status, or APOE genotype. The association likely reflects the well-documented enrichment of functional regulatory variation on African-ancestry haplotypes: greater sequence diversity on YRI-derived segments translates into more heterozygous sites, more opportunities to harbor cis-regulatory variants, and consequently a higher per-individual rate of detected allelic imbalance (Stranger et al., 2007; Aguet et al., 2017). The comparatively weak contribution of PEL ancestry to ASE burden is consistent with the narrower spread of Amerindian ancestry proportions within this cohort and the reduced power this confers for detecting ancestry-linked regulatory effects at the global level, rather than an absence of Amerindian regulatory variation per se.

Two non-mutually exclusive biological mechanisms could explain ancestry-associated ASE differences and distinguishing them is central to interpreting the ASED results and the directional analysis that follows. First, ancestry-associated cis-regulatory variants may differ in frequency or effect size across populations, such that haplotypes from one ancestry carry a higher density of variants that drive allelic imbalance (variant frequency mechanism). Second, the regulatory environment, including chromatin state, transcription factor occupancy, and long-range interactions, may differ between ancestral haplotypes in ways that are not captured by single variant effects (regulatory context mechanism). The ASED results primarily test the first mechanism across many genes. The ancestry-ASE scan directly tests the second by conditioning on within-individual haplotype ancestry at each locus. The directional analysis distinguishes which ancestral regulatory program is dominant when haplotypes diverge.

Gene-level population comparisons using the ASED framework identified substantial heterogeneity in allelic imbalance across groups, with the strongest contrasts observed between African American and non-Hispanic White individuals. The presence of ASED signals at regions of known population-differentiated regulatory architecture, including the MHC, is consistent with group-level ASE differences reflecting genuine ancestry-linked variation in cis-regulatory mechanisms. A notable feature of the ASED results is that a substantial proportion of significant genes showed opposing imbalance directions across population groups, suggesting that the same locus can harbor distinct regulatory architectures on different ancestral haplotype backgrounds. This pattern is conceptually distinct from uniform expression differences across groups and underscores the value of ancestry-aware analysis for disentangling the sources of regulatory variation. The observation of group differences in allelic imbalance at specific genes is consistent with several non-mutually exclusive mechanisms. First, the two groups may carry different frequencies of cis-regulatory variants that directly alter (Mogil et al., 2018) transcription (Lam et al., 2017) factor binding or splicing. Second, the groups may differ in haplotype background, such that variants in linkage disequilibrium with functional cis-regulatory elements drive the observed imbalance rather than the tested variant itself. Third, opposing effects may exist within a group such that different individuals within the same population may show allelic imbalance in opposite directions, producing apparent between-group differences in the mean that are driven by within-group heterogeneity in the direction of effect, which is the pattern expected from ordinary ASE at a subset of genes. The difference is that such imbalance would be distributed randomly within and across groups rather than tracking ancestry. This third scenario is formally distinct from ASE that is completely independent of ancestry, because it predicts a specific relationship between ancestry frequency and the direction of imbalance at the individual level. The directional analysis described above was designed to distinguish these scenarios: a consistent directional signal across individuals indicates a fixed regulatory architecture, whereas a mixed or balanced signal indicates either no ancestry effect or opposing within-group effects that “cancel” at the population level.

Ancestry-ASE scan recapitulates known regions of population-differentiated regulatory architecture and extends them to a multi-ancestry admixed cohort. Only one locus, chr6:32–33 Mb, showed calls in opposite directions under different baselines, and these calls carry a mapping bias flag, so the evidence that the direction of regulatory effect depends on ancestral background, rather than only its magnitude, remains limited. Future work examining whether the 52 ancestry-associated ASE loci identified here are enriched for population-differentiated regulatory elements, such as ancestry-specific eQTLs, differentially methylated regions, or population-stratified chromatin accessibility peaks, would help establish the molecular basis of these signals.

The ancestry-directional analysis adds a dimension that the ancestry-ASE scan alone cannot provide, not just whether admixed configurations alter the rate of ASE, but which ancestral haplotype is preferentially expressed when they do. The global symmetry of the directional signal, mean log_2_ ancestry ratios indistinguishable from zero across all baselines and populations, rules out a scenario in which one ancestral regulatory architecture globally dominates the admixed transcriptome, and grounds the locus-specific directional findings as genuine exceptions. The identification of the *SLC24A5* locus as a CEU-dominant positive control validates the haplotype-resolved methodology. Among the novel directional findings, the *CSMD1* locus illustrates the biological insight uniquely accessible through this framework: consistent non-European haplotype dominance across populations with distinct non-European ancestries points to a European-specific cis-regulatory architecture that is subordinate when paired with either African or Amerindian haplotypes. This pattern was consistent across baselines, unflagged for reference bias, and statistically robust (minimum q = 8.92 × 10^−99^), which identifies *CSMD1* as a candidate for follow-up functional characterization in the context of its well-documented associations with schizophrenia, bipolar disorder, and cognitive traits in genome-wide association studies (Ripke et al., 2011; Havik et al., 2011; Koiliari et al., 2014). The absence of *CSMD1* from the gene-level analysis reflects a distinction between the two approaches: the gene-level analysis tests whether individual genes within a significant window reach significance for admixture-associated ASE, whereas the directional analysis detects consistent haplotype-expression bias aggregated across all sites within a window. A locus can show strong directional bias without any single gene reaching gene-level significance, particularly when the effect is distributed across multiple genes or non-coding elements within the window.

Taken together, the three analytical components of this study capture complementary dimensions of ancestry-dependent ASE. Global ancestry modeling shows that individuals with greater YRI ancestry proportion carry more ASE-positive genes overall (R² = 0.194 for protein-coding ASE genes in the fully adjusted model, incremental R² = 0.164 above covariates alone), indicating that African ancestry is associated with increased ASE frequency across the transcriptome. The ancestry-ASE scan identifies loci where LAZ-heterozygous configurations are associated with elevated ASE, and the directional analysis reveals which ancestral haplotype is preferentially expressed at those loci. The direction of haplotype-specific expression is population-specific and locus-specific rather than globally fixed and reflects local cis-regulatory architecture at each gene. The most parsimonious interpretation is that African haplotypes more frequently carry cis-regulatory variants that diverge from European haplotypes, generating allelic imbalance when the two are paired in an admixed individual, and that the direction of that imbalance is determined by the local regulatory environment rather than a genome-wide dominance of one ancestral regulatory program. The prominence of the African ancestry signal likely reflects the greater genetic diversity of African-descent haplotypes relative to European-descent haplotypes, a well-established consequence of the out-of-Africa bottleneck (Tishkoff et al., 2009; Campbell & Tishkoff, 2008). African haplotypes carry a higher density of common regulatory variants, a broader spectrum of cis-regulatory alleles, and more population-private variants at regulatory sites (Stranger et al., 2007; Lappalainen et al., 2013; Campbell & Tishkoff, 2008), all of which increase the probability of cis-regulatory divergence when paired with a European haplotype in an admixed individual. Future work integrating phased long-read sequencing or chromosome-level haplotype reconstruction could enable direct parental haplotype assignment in admixed individuals, allowing finer-grained characterization of the regulatory architecture underlying these ancestry-ASE signals (Stranger et al., 2007; Lappalainen et al., 2013; Campbell & Tishkoff, 2008)

Several features of the study design are worth considering when interpreting these findings. The cohort consists of older adults, a context in which regulatory variation and its downstream consequences may differ from younger reference populations. Although clinical and demographic covariates did not substantially alter the ancestry associations observed here, the aged whole-blood context may modulate the magnitude of ancestry effects at specific loci (Martinez-Jimenez et al., 2017), particularly those with known immune or inflammatory regulatory functions. Whether the ancestry-associated ASE patterns identified here generalize across tissues or are amplified by age-related transcriptional changes remains an open question.

This study has several limitations. First, analyses were performed in whole blood, and cis-regulatory architectures are tissue and cell-type specific, so the findings may not extend to other tissues. Second, whole blood is a heterogeneous tissue, and inter-individual variation in cell-type composition, which itself varies by ancestry (Randolph et al., 2021; Aran et al., 2017), represents a potential confound. If individuals of African ancestry carry a systematically different distribution of blood cell types relative to individuals of European ancestry, and if those cell types have distinct cis-regulatory architectures, the observed ancestry-associated ASE signals could partially reflect cell-type-specific regulatory variation rather than ancestry-specific cis-regulatory effects. Deconvolution of bulk RNA-seq into cell-type-specific expression profiles, or replication in sorted cell populations, would be required to fully distinguish these contributions. Third, local ancestry inference introduces classification uncertainty at locus boundaries and in regions of high linkage disequilibrium, which may attenuate ancestry-ASE scan signals at some loci (Browning et al., 2023). The frequency and genomic distribution of within-window ancestry configuration switching, which reflects this uncertainty, are characterized in Figures S3 and S4. Fourth, the GRCh38 reference genome was constructed from a sample of limited ancestral diversity, which may introduce mapping bias at variants where the reference allele is enriched in European-ancestry populations. We have flagged potentially affected windows using a reference-bias indicator in the supplementary tables. Fifth, allele frequencies at heterozygous sites differ across ancestries independently of regulatory function, which can produce apparent imbalance through ascertainment, and our analyses do not separate this effect from differences in regulatory architecture. Sixth, the ancestry combinations observed in each population group depend on who was sampled, and some combinations may be over- or under-represented relative to the wider population. Additionally, the power to detect ancestry-associated ASE at any given locus is constrained by the frequency of LAZ-heterozygous configurations at that locus within the cohort. Regions where admixed individuals rarely carry two different ancestries will be systematically underpowered regardless of sample size. Finally, the cross-population replication criterion, while powerful as a significance filter, is conservative and is additionally constrained by the frequency of LAZ-heterozygous configurations at each locus.

Regions where few individuals carry two different ancestral haplotypes will have limited power regardless of sample size. True loci with ancestry-specific effects limited to a single population will be missed by design. Population-specific regulatory effects could be explored in future work by examining loci reaching significance in a single admixed population prior to cross-population filtering, though interpretation of such signals would require careful consideration of population-specific sample sizes and the elevated risk of false positives in the absence of independent replication.

A further consideration is the potential contribution of genomic imprinting to the observed ancestry-associated ASE signals. Imprint control regions (ICRs) regulate parent-of-origin-specific gene expression through differential DNA methylation established before tissue specification, and ancestry-associated differences in haplotype-specific expression at imprinted loci could reflect imprinting rather than ancestry-linked cis-regulation. To assess this, we cross-referenced our results against the 1,488 candidate ICRs from the human imprintome (Jima et al., 2022). Twelve of the 33 replicated CEU/CEU scan windows (Table 3) contained at least one candidate ICR. However, none of the significant ancestry-associated genes within those windows were proximal to the overlapping ICR, suggesting the window-level overlap is incidental given the 1 Mb window size. Among the 1,537 ASED-significant genes, 59 (3.8%) overlapped a candidate ICR, compared to 2.9% of all tested genes (OR = 1.42, p = 0.019). *KCNQ1*, a well-characterized maternally imprinted gene, was among the ASED-significant genes overlapping ICRs and should be interpreted with caution. Taken together, imprinting is unlikely to account for the majority of the ancestry-associated ASE signals identified here, though it cannot be fully excluded for individual genes without parent-of-origin phasing data.

Despite these limitations, the findings have substantive implications for the study of regulatory variation in diverse populations. The demonstration that LAZ predicts ASE at dozens of loci and that the direction of ancestry effects is locus-specific rather than globally biased supports a model in which ancestry-dependent cis-regulatory variation is a widespread feature of the admixed transcriptome, distributed across the genome and reflective of population-differentiated regulatory architecture at discrete loci. This has practical consequences for gene expression prediction models: frameworks trained on homogeneous European reference panels will systematically misrepresent the regulatory state at loci identified by the ancestry-ASE scan.

Incorporating local ancestry as a locus-level predictor in expression modeling pipelines is a tractable next step that this study motivates. More broadly, the results reinforce the case for ancestry-diverse study designs in regulatory genomics: the signals identified here were accessible precisely because admixture generates within-individual contrasts across haplotype backgrounds, a natural experiment unavailable in homogeneous cohorts.

In summary, we present a multi-scale characterization of ancestry effects on allele-specific expression in a cohort of admixed older adults. Global ASE burden, gene-level population comparisons, and a genome-wide ancestry-configuration scan demonstrate a consistent picture: genetic ancestry, particularly African-ancestry haplotypes, is a major source of cis-regulatory variation, and this relationship manifests at both the genome-wide level and at discrete loci with ancestry-specific regulatory architectures. The ancestry-ASE scan framework provides a principled and replication-controlled approach to identifying these loci, and the complementary directional analysis reveals the direction of ancestry effects, specifically which ancestral haplotype dominates expression and whether that dominance is consistent across populations with different admixture backgrounds, together establishing a framework for ancestry-aware characterization of cis-regulatory variation in admixed populations.

## Supporting information

Supplementary Technical Methods

## Acknowledgements

This work was supported by the National Institute on Aging (R01AG070935).

---

**Figure S1.**
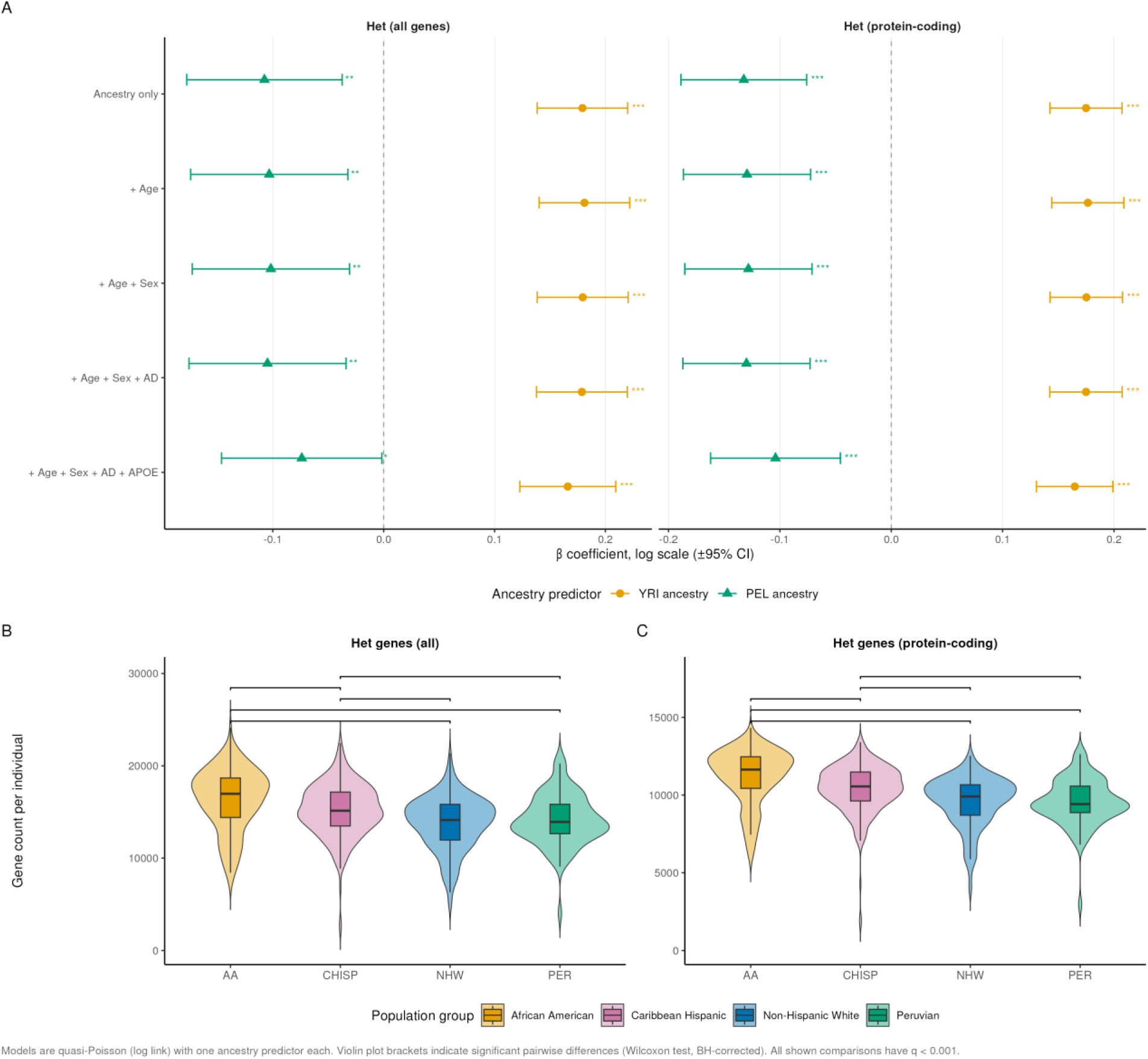
Ancestry predicts heterozygous gene burden across population groups. (A) Beta coefficients (±95% CI) from quasi-Poisson generalized linear models predicting heterozygous gene counts (all genes and protein-coding genes) as a function of YRI ancestry proportion (orange circles) or PEL ancestry proportion (green triangles), across five covariate adjustment sets. Asterisks denote significance: *p<0.05, **p<0.01, ***p<0.001. (B, C) Distribution of per-individual heterozygous gene counts by population group for all genes (B) and protein-coding genes (C). Brackets indicate significant pairwise differences by Wilcoxon rank-sum test with Benjamini-Hochberg correction. All shown comparisons have q<0.001. AA, African American; CHISP, Caribbean Hispanic; NHW, Non-Hispanic White; PER, Peruvian.

**Figure S2.**
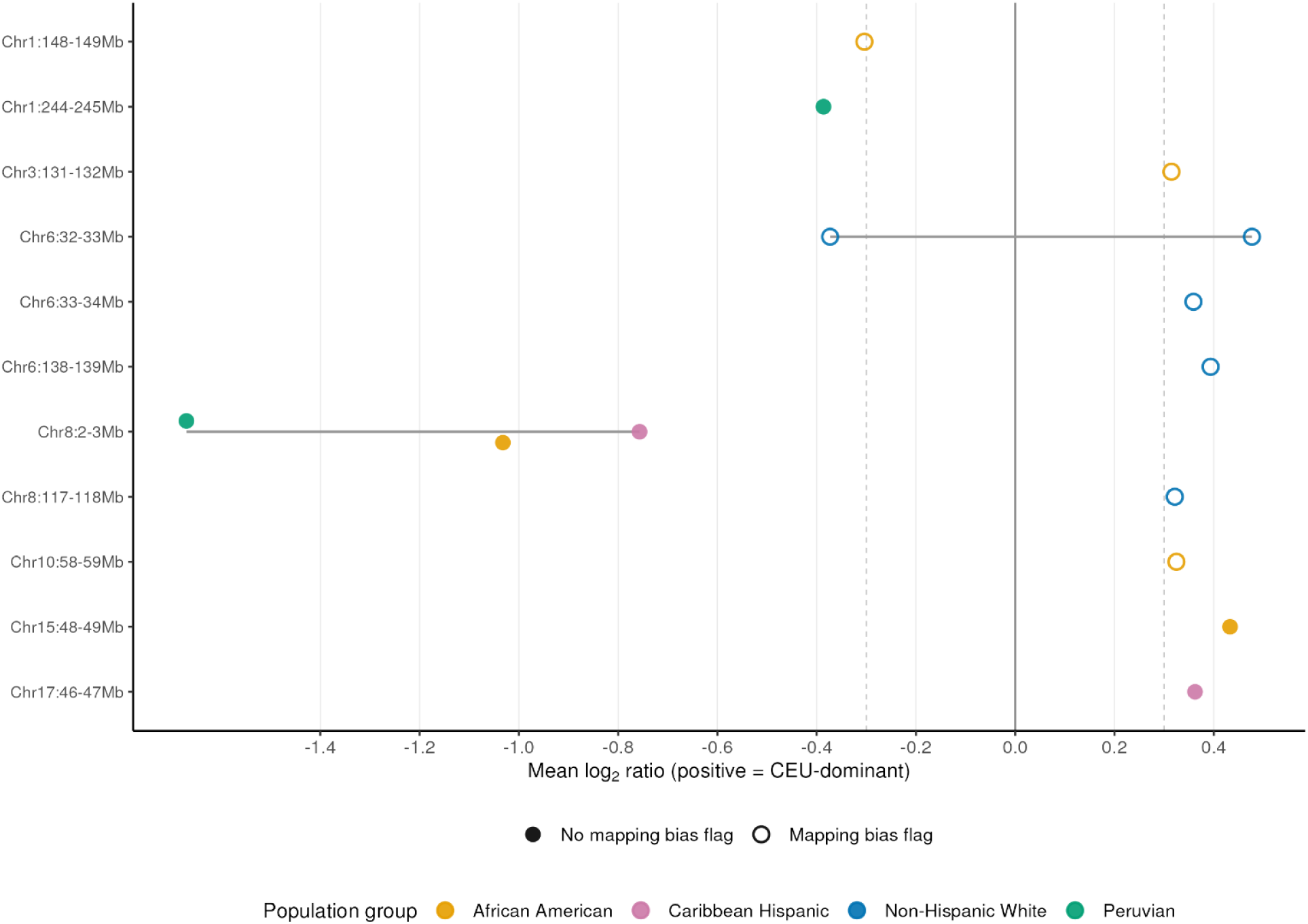
Directional analysis of ancestry-associated ASE at high-confidence loci. Lollipop plot showing the mean log2 ratio of CEU-dominant to non-CEU-dominant allele-specific expression per locus and population group for the 13 high-confidence directional calls at 11 loci in Supplementary Table S5. Positive values indicate CEU haplotype dominance and negative values indicate YRI or PEL haplotype dominance. Dashed lines mark the classification threshold of ±0.3. Filled points have no mapping bias flag and open points have a mapping bias flag. The grey horizontal segment per locus spans the range across points. A locus can have calls in more than one direction under different baselines, as at Chr6:32–33 Mb. Chr8:2–3 Mb is the CSMD1 locus and the only locus with the same direction (PEL-dominant) in at least two population groups. AA, African American; CHISP, Caribbean Hispanic; NHW, Non-Hispanic White; PER, Peruvian.

**Figure S3.**
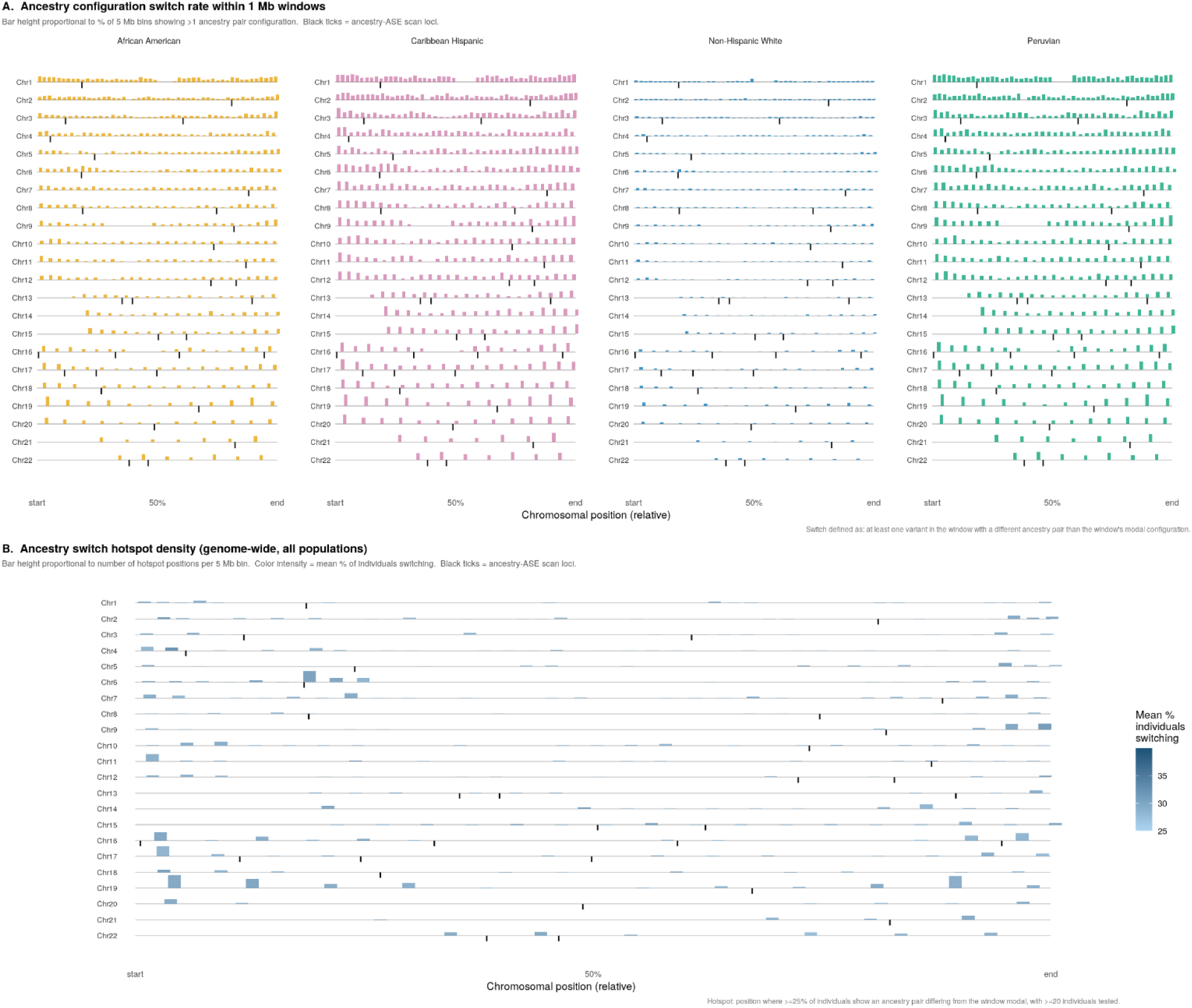
Ancestry configuration switching within 1 Mb windows across the genome. (A) Ancestry configuration switch rate by population. Each panel shows one of the four MAGENTA cohort populations (African American, Caribbean Hispanic, Non-Hispanic White, Peruvian). Chromosomes 1–22 are displayed as horizontal tracks. Ancestry switching was assessed within 1 Mb analysis windows; for display, windows were aggregated into 5 Mb bins and bar height reflects the percentage of bins in which at least one 1 Mb window showed a switch. Ancestry pairs were assigned in a haplotype-dependent manner, such that CEU/YRI and YRI/CEU represent distinct configurations reflecting which haplotype carries which ancestral origin. A switch was defined as a window in which at least one variant carried a local ancestry pair differing from the most common configuration observed across all variants in that window for that individual, indicating that a local ancestry tract boundary fell within the window. For example, if 28 of 30 variants in a window carried a CEU/YRI ancestry pair but 2 variants carried a YRI/CEU pair, that window was counted as a switch for that individual. Black tick marks indicate the positions of the 33 ancestry-ASE scan loci identified in the main analysis. Bar height reflects bin-level detection frequency and is influenced by sample size; populations with smaller sample sizes (Peruvian, n=83) may show elevated bar heights relative to their population-level switch rate. Per-window switch rates, which account for sample size, are the more appropriate basis for cross-population comparison: 6.7% in Non-Hispanic White individuals (reflecting local ancestry inference uncertainty rather than true admixture), 21.1% in African Americans, 34.1% in Peruvians, and 37.8% in Caribbean Hispanic individuals. **(B) Genome-wide ancestry switch hotspot density (all populations combined).** Bar height reflects the number of hotspot positions per 5 Mb bin across all populations. A hotspot position was defined as a genomic site at which at least 25% of individuals tested at that position carried an ancestry pair differing from the modal configuration of their 1 Mb window, with a minimum of 20 individuals tested. Color intensity reflects the mean percentage of individuals switching at hotspot positions within each bin (scale at right). Black tick marks indicate the 33 ancestry-ASE scan loci. Hotspots are distributed across the genome with a notable concentration on Chr6, consistent with the high recombination rate and complex ancestry structure of the major histocompatibility complex region. The absence of hotspot enrichment at the majority of ancestry-ASE scan loci supports the interpretation that identified loci reflect stable ancestry-associated regulatory effects rather than local ancestry inference artifacts.

**Figure S4.**
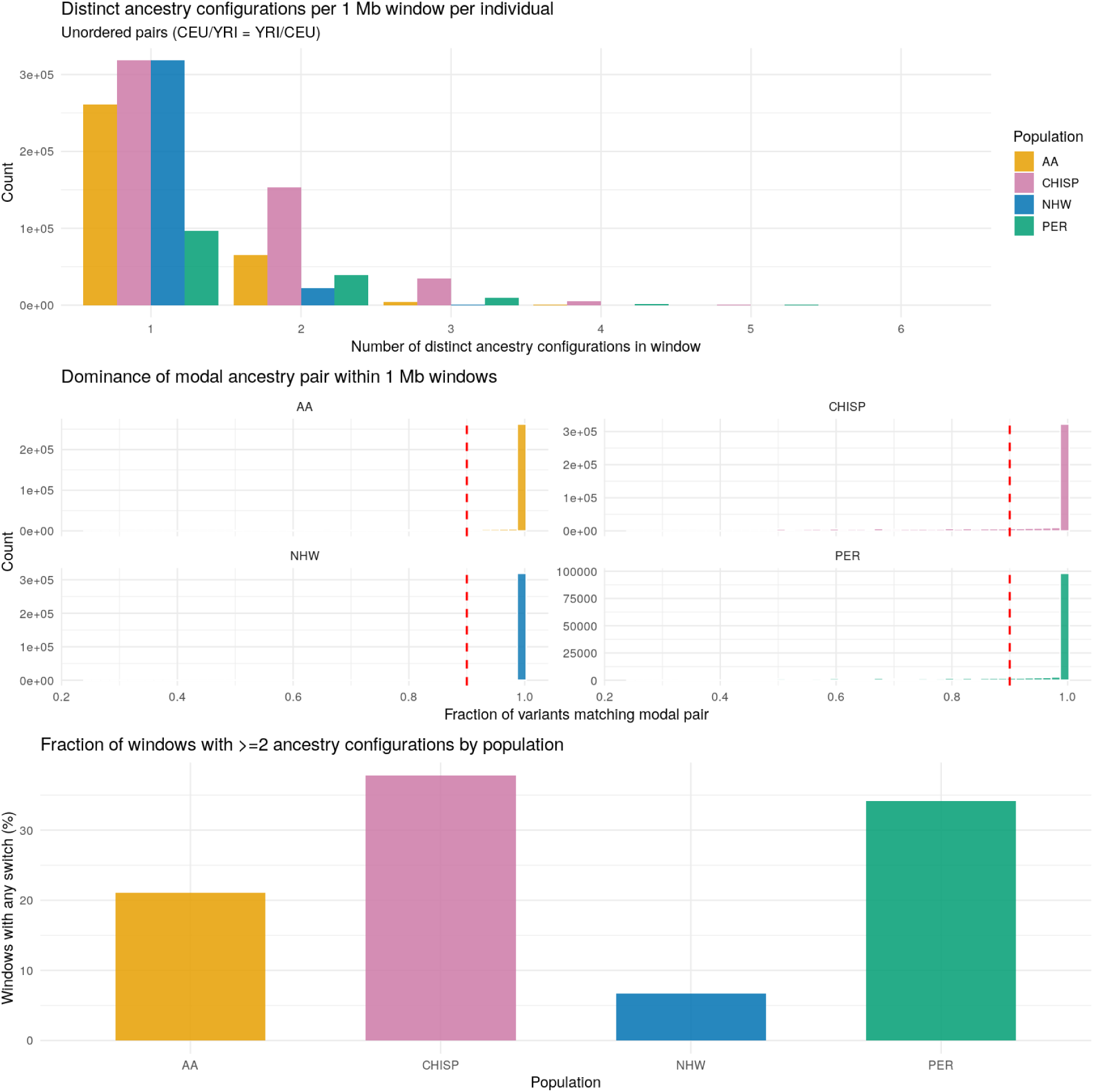
Characterization of local ancestry configuration switching within 1 Mb windows across the genome. (A) Distribution of distinct ancestry configurations per window. Bars show the number of window x individual observations with 1, 2, 3, or more distinct unordered ancestry pair configurations within a single 1 Mb window, by population. Ancestry pairs are treated as unordered (CEU/YRI = YRI/CEU). The majority of windows show a single configuration, with a rapid drop-off at 2 or more. **(B) Dominance of the modal ancestry pair within windows.** For each per-individual per-window observation, the modal pair fraction was calculated as the proportion of variants within the window carrying the most common ancestry pair configuration for that individual. A modal pair fraction of 1.0 indicates a fully homogeneous window where every variant carries the same ancestry pair. The red dashed line marks 0.90, the threshold used to define windows with a dominant ancestry configuration. Across all populations, 82.4% of windows met this threshold. Non-Hispanic White and African American individuals show distributions concentrated almost entirely at 1.0, consistent with low admixture complexity and longer ancestry tracts relative to window size. Caribbean Hispanic and Peruvian individuals show left-shifted tails, reflecting more frequent within-window ancestry transitions driven by three-way admixture and greater tract fragmentation. **(C) Per-population switch rate.** Bars show the percentage of windows per individual with at least one variant carrying an ancestry pair differing from the window modal configuration: 6.7% in Non-Hispanic White individuals, 21.1% in African Americans, 34.1% in Peruvians, and 37.8% in Caribbean Hispanic individuals. The low rate in Non-Hispanic White individuals, who are predominantly CEU/CEU homozygous, reflects local ancestry inference uncertainty rather than true admixture and establishes a noise floor for the analysis.

## Notes

### Competing Interest Statement

The authors have declared no competing interest.

