## Supplementary Technical Methods for "Genetic ancestry influences the relative expression of ancestral haplotypes within admixed individuals"

**Ancestry-ASE Scan: Complete Statistical Specification**

*Vazquez-Hidalgo et al.*

### **1. Overview and Design Rationale**

The ancestry-ASE scan was designed to identify genomic loci at which the local ancestry configuration of an individual's two haplotypes is systematically associated with the probability of allele-specific expression (ASE) at heterozygous variants within that locus. The central question is whether individuals carrying one or two copies of a particular ancestral haplotype show a different propensity for allelic regulatory imbalance compared to individuals carrying two copies of a reference ancestry.

The scan takes advantage of naturally occurring variation in local ancestry in admixed populations. In an admixed individual, each autosomal locus carries haplotypes from two potentially distinct ancestral backgrounds. When the cis-regulatory architecture differs between ancestral populations at a given locus, this difference manifests as a higher rate of ASE in individuals carrying one copy of each ancestry than in individuals carrying two copies of the same ancestry. The scan tests this prediction using a windowed logistic regression. Within each 1 Mb genomic window, every heterozygous ASE-informative variant in every individual contributes an observation. The binary outcome is ASE status and the predictor is the local ancestry pair configuration at that variant, with a homozygous ancestry serving as the reference category.

### **2. Input Data and Preprocessing**

#### **2.1 Allelic Read Counts**

Input data consist of per-variant allelic read counts generated by phASER (Castel et al., 2016). For each individual i and heterozygous variant j, the inputs are reference allele count r_ij, alternative allele count a_ij, and total reads n_ij = r_ij + a_ij, along with the local ancestry assignment for each haplotype from FLARE (Browning et al., 2023).

#### **2.2 Quality Filters**

Sites were retained if: (1) n_ij >= 20; (2) local ancestry was assigned to both haplotypes; (3) haplotype labels were valid (1 or 2); (4) sample ID, variant ID, position, allele, and read counts were all non-missing.

#### **2.3 Ancestry Pair Derivation**

For each variant and individual, let anc_h1 and anc_h2 be the local ancestry of haplotypes 1 and 2. The ancestry pair was defined as paste(min(anc_h1, anc_h2), max(anc_h1, anc_h2), sep='/'), ensuring CEU/YRI and YRI/CEU are treated identically.

### **3. ASE Calling**

#### **3.1 Binomial Test**

Under the null of equal allelic expression (p = 0.5), the two-sided p-value at site j for individual i is:

p_raw = 2 x pbinom(min(r_ij, a_ij), size = n_ij, prob = 0.5)

Values exceeding 1 are capped at 1. P-values are adjusted genome-wide within each individual using the Benjamini-Hochberg procedure.

#### **3.2 Effect Size and Classification**

The log2 allelic fold change is:

log2AFC_ij = log2[(r_ij + 0.5) / (a_ij + 0.5)]

A site is classified as ASE-positive (ASE_ij = 1) if q <= 0.05 AND |log2AFC| >= 1.0. The dual threshold excludes high-coverage sites with trivial imbalance and unreliable estimates at low coverage.

### **4. Windowed Logistic Regression (Row Scan)**

#### **4.1 Window Construction**

window_start = floor((pos - 1) / 10^6) x 10^6 + 1; window_end = window_start + 10^6 - 1

Non-overlapping 1 Mb windows in 1-based coordinates. Windows with fewer than 200 total observations were excluded.

#### **4.2 Model Specification**

Within window w and population p, the logistic regression model is:

logit[Pr(ASE_ij = 1)] = B0 + sum_k Bk * I[AncestryPair_ij = k]

Each observation is one variant j in one individual i (row-level data; n approximately 3,000 to 8,000 per window), and the sum is over non-reference ancestry pair levels. The intercept B0 represents the log-odds of ASE in the baseline (homozygous) population. Each Bk is the log odds ratio for ancestry pair k versus the baseline. The odds ratio is OR = exp(Bk) and the 95% CI is [exp(Bk +/- 1.96*SE_k)]. Models were fit by maximum likelihood via R's glm(family = binomial). Non-converging models were discarded.

#### **4.3 Baseline Specification and Multiple Testing**

Three independent scans were run with baselines CEU/CEU, YRI/YRI, and PEL/PEL. Within each population, p-values across all windows and terms were adjusted genome-wide by Benjamini-Hochberg FDR. Genomic inflation lambda = 3.32 reflects correlated observations within windows (multiple variants per individual are not independent) rather than model misspecification.

### **5. Cross-Population Replication and Locus Definition**

#### **5.1 Replication Criterion**

A window was retained as significant if it reached q < 0.05 in at least 2 of 4 populations (AA, CHISP, NHW, PER) for the same term within the same baseline. Under the null, the probability of a false positive across four independent tests is alpha^4 = 0.05^4 = 6.25 x 10^-6, providing stringent family-wise error rate control despite genomic inflation within individual scans. This filter reduced 447 nominally significant windows to 33, 11, and 16 replicated loci for the CEU/CEU, YRI/YRI, and PEL/PEL baselines, respectively.

#### **5.2 Locus Clustering**

Significant windows were clustered by genomic overlap. Windows were sorted by chromosomal position and merged greedily when adjacent windows overlapped. Each cluster received a unique locus identifier. The lead hit per locus per population was the window with minimum q-value (ties broken by maximum |z|).

### **6. COLLAPSED Scan (Validation)**

In the collapsed scan, data were aggregated to the per-individual per-window level: k_iw = number of ASE-positive variants and m_iw = total variants for individual i in window w. The model glm(cbind(k, m-k) ~ ancestry_pair, family=binomial) eliminates within-individual correlation. Results correlated at r = 0.9999 with the row scan (lambda = 3.32 in both), confirming the row scan signal is not an artifact of variant-level pseudoreplication.

### **7. Ancestry-Associated Gene Analysis**

The window-level logistic model was refit at the gene level for all genes overlapping a significant locus, using only variants within that gene (j in gene g). P-values were adjusted within each baseline by Benjamini-Hochberg FDR. Genes reaching q < 0.05 were classified as ancestry-associated genes. The ancestry-associated gene reported for each locus is the gene with the lowest q-value within the locus boundaries. Gene annotations were drawn from GENCODE v43 (GRCh38); gene biotype from the gene_type GTF attribute.

### **8. Ancestry-Directional ASE Analysis**

#### **8.1 Haplotype-Resolved Expression Ratio**

For each site within a significant locus, let C_anc1,ij and C_anc2,ij be allelic read counts assigned by local ancestry rather than by reference/alternative identity, which is reference-genome-dependent and can confound ancestry assignment. The per-site log2 ancestry ratio is:

R_ij = log2[(C_anc1,ij + 1) / (C_anc2,ij + 1)]

Sites with C_anc1 + C_anc2 < 10 were excluded. Only sites with exactly one haplotype of each target ancestry were included.

#### **8.2 Site Classification**

anc1_dominant: R_ij > 0.5; anc2_dominant: R_ij < -0.5; balanced: |R_ij| <= 0.5

#### **8.3 Window-Level Test**

Let n_anc1 and n_anc2 be the counts of anc1- and anc2-dominant sites within a window and population. A two-sided binomial test under H0: p = 0.5:

p_binom = 2 x B(min(n_anc1, n_anc2); n_anc1 + n_anc2, 0.5)

A window is classified as anc1_dominant if the mean log2 ratio R-bar_w > 0.3 AND p_binom < 0.05; anc2_dominant if R-bar_w < -0.3 AND p_binom < 0.05; and balanced_or_mixed otherwise. FDR correction was applied by Benjamini-Hochberg within each baseline. Reliability filters required n_samples >= 10 and n_sites >= 50.

#### **8.4 Mapping Bias Flag**

Let f_ref be the proportion of directional calls where the dominant allele is the GRCh38 reference allele. The flag is raised if f_ref > 0.70 or f_ref < 0.30. Because GRCh38 was constructed predominantly from European-ancestry sequence, a high f_ref at CEU-dominant loci is consistent with a reference allele mapping advantage rather than true regulatory dominance.

### **9. Parameter Summary**

| **Parameter** | **Value** | **Code variable** | **Rationale** |
| --- | --- | --- | --- |
| MIN_TOTAL_READS | 20 | MIN_TOTAL_READS | Minimum read depth for reliable binomial ASE testing |
| ASE_Q | 0.05 | ASE_Q | FDR threshold for within-individual ASE calling |
| ASE_MIN_ABS_LOG2AFC | 1.0 | ASE_MIN_ABS_LOG2AFC | ≥2-fold allelic imbalance required; prevents trivial significance |
| WINDOW_BP | 1,000,000 | WINDOW_BP | 1 Mb windows balance power and resolution |
| MIN_N_PER_WINDOW | 200 | MIN_N_PER_WINDOW | Minimum observations for stable GLM fitting |
| CROSSPOP_SHARED_MIN_POPS | 2 | CROSSPOP_SHARED_MIN_POPS | Requires replication in ≥2 populations; P ≈1.4x10⁻⁵ under null |
| PSEUDOCOUNT (directional) | 1 | PSEUDOCOUNT | Add-one smoothing for log2 ratio at low counts |
| MIN_READS (directional) | 10 | MIN_READS | Minimum ancestry-assigned reads for directional analysis |
| Directional \|R-bar\| threshold | 0.3 | mean_log2_ratio threshold | Requires moderate window-level directionality in addition to binomial p |
| Mapping bias flag | >0.70 or <0.30 | mapping_bias_flag | Flags potential GRCh38 reference allele alignment advantage |
| Min samples (directional) | 10 | n_samples >= 10 | Minimum individuals with directional data per window |
| Min sites (directional) | 50 | n_sites >= 50 | Minimum informative sites for robust binomial testing |

### **10. Software and Reproducibility**

All analyses were performed in R version 4.3.1 (2023-06-16). Scan: data.table, base R glm(). Directional analysis: data.table, parallel::mclapply() (4 cores). Local ancestry inference: FLARE (Browning et al., 2023), 1000 Genomes Project high-coverage reference panels (Byrska-Bishop et al., 2022). ASE quantification: phASER (Castel et al., 2016). Gene annotations: GENCODE v43 (GRCh38). All coordinates GRCh38 (hg38). Analysis code is available at [GitHub URL].

### **11. Conceptual Overview of the Analytical Approach**

This section describes the analytical logic of the ancestry-ASE scan in non-technical terms. The formal statistical specification is given in Sections 1 through 9 above.

#### **Admixture as a natural experiment**

An admixed individual, for example a person of African American ancestry, carries one copy of each chromosome from each biological parent. Because their ancestors came from genetically distinct populations, different segments of each chromosome trace back to different ancestral populations. At any given genomic region, an admixed person may carry one chromosome segment of African ancestry and one of European ancestry, or two segments of the same ancestry. This configuration is described by local ancestry zygosity.

If two ancestral populations differ in how they regulate a gene at a given locus, perhaps because they carry different cis-regulatory variants at different frequencies, then a person carrying one haplotype from each ancestry will show more allelic imbalance at that locus than a person carrying two haplotypes from the same ancestry. When two haplotypes carry similar regulatory programs, allelic expression tends to be balanced. When they carry divergent programs, that divergence appears as allele-specific expression.

#### **Calling ASE at individual variants**

For each heterozygous variant in each individual, we count how many RNA-seq reads come from the reference allele and how many from the alternative allele. Under the null hypothesis of equal allelic expression, these counts should follow a binomial distribution with p = 0.5. We apply a binomial test to determine whether the observed imbalance differs significantly from 50/50, and also require the imbalance to be at least 2-fold to exclude statistically significant but biologically trivial imbalances at very high-coverage sites. Sites passing both thresholds are classified as ASE-positive.

#### **Aggregating to genomic windows**

We divide the genome into non-overlapping 1 Mb windows. Within each window, we pool all heterozygous ASE-informative variant observations across all individuals. Each observation records whether that variant showed ASE in that individual and what ancestry pair that individual carries at that locus, derived from local ancestry inference by FLARE. Possible ancestry pairs include, for example, CEU/CEU, CEU/YRI, or YRI/YRI.

#### **The logistic regression**

We fit a logistic regression in each window with ASE status (0 or 1) as the outcome and ancestry pair as the categorical predictor. The reference category is the homozygous ancestry of interest, for example CEU/CEU in the CEU baseline scan. The model estimates, for each non-reference ancestry pair, the log odds ratio of ASE relative to the homozygous baseline. A significant positive coefficient for the heterozygous pair, such as CEU/YRI, indicates that individuals carrying one CEU and one YRI haplotype in that window have a higher rate of ASE than individuals carrying two CEU haplotypes, consistent with divergent regulatory architecture between the two ancestries at that locus.

Because each individual contributes many variant observations per window, the observations within a window are not fully independent. This produces an elevated test statistic (lambda = 3.32). Rather than applying genomic control, we address this through cross-population replication: a genuine ancestry-ASE signal should appear across multiple independent population groups, while inflation-driven false positives are unlikely to replicate.

#### **Cross-population replication**

We run three separate scans, one for each homozygous baseline (CEU/CEU, YRI/YRI, PEL/PEL). A window is retained only if it reaches FDR < 5% in at least two of the four population groups (NHW, AA, CHISP, PER) within the same baseline scan. Because each population group is genetically and demographically distinct, a false positive driven by population-specific confounding is unlikely to appear in more than one group. Under the null, the probability of a false positive in two or more independent groups at alpha = 0.05 is approximately 1.4 x 10^-5. Replicated windows that overlap genomically are merged into non-redundant loci.

#### **Identifying ancestry-associated genes**

A significant window may span many genes. To identify which gene is driving the ancestry-ASE association, we refit the logistic regression model at the gene level, restricting observations to variants within each gene's boundaries. Genes reaching FDR < 5% within a significant locus are classified as ancestry-associated genes. The gene with the lowest q-value within the locus is reported as the primary ancestry-associated gene for that locus.

#### **Directional analysis**

The logistic regression indicates that a locus shows differential ASE by ancestry but does not specify which ancestral haplotype tends to be expressed at higher levels. To address this, we use the phased structure of the data. Because we know which allele at each variant came from which haplotype, and because FLARE assigns each haplotype to an ancestral population, we can determine for each heterozygous site whether the allele from ancestry 1 or ancestry 2 is more highly expressed. We summarize this across all sites within a window using the log2 ancestry ratio R_ij and test whether the window shows a consistent directional bias using a binomial test. A window is classified as ancestry-dominant when one haplotype's allele is systematically expressed at higher levels, and as balanced or mixed otherwise. We flag windows where the dominant allele frequently matches the GRCh38 reference allele, since GRCh38 was constructed predominantly from European-ancestry sequence and reference allele mapping bias could inflate apparent CEU dominance.
